# The yeast RNA methylation complex consists of conserved yet reconfigured components with m6A-dependent and independent roles

**DOI:** 10.1101/2023.02.10.528004

**Authors:** Imke Ensinck, Alexander Maman, Waleed S. Albihlal, Michelangelo Lassandro, Giulia Salzano, Theodora Sideri, Steven Howell, Enrica Calvani, Harshil Patel, G. Guy Bushkin, Markus Ralser, Ambrosius P. Snijders, Mark Skehel, Ana Casañal, Schraga Schwartz, Folkert J. van Werven

**Affiliations:** The Francis Crick Institute, 1 Midland Road, London, NW1 1AT, UK; Department of Molecular Genetics, Weizmann Institute of Science, Rehovot, Israel; Charité Universitätsmedizin Berlin, Department of Biochemistry, Germany; Human Technopole, Viale Rita Levi-Montalcini, 1, 20157 Milan, Italy; Whitehead Institute for Biomedical Research, Cambridge, MA 02142, USA

## Abstract

*N6*-methyladenosine (m6A), the most abundant mRNA modification, is deposited in mammals/insects/plants by m6A methyltransferase complexes (MTC) comprising a catalytic subunit and at least five additional proteins. The yeast MTC is critical for meiosis and was known to comprise three proteins, of which two were conserved. We uncover three novel MTC components (Kar4/Ygl036w-Vir1/Dyn2). All MTC subunits, except for Dyn2, are essential for m6A deposition and have corresponding mammalian MTC orthologs. Unlike the mammalian bipartite MTC, the yeast MTC is unipartite, yet multifunctional. The mRNA interacting module, comprising Ime4, Mum2, Vir1, and Kar4, exerts the MTC’s m6A-independent function, while Slz1 enables the MTC catalytic function in m6A deposition. Both functions are critical for meiotic progression. Kar4 also has a mechanistically separate role from the MTC during mating. The yeast MTC constituents play distinguishable m6A-dependent, MTC-dependent and MTC-independent functions, highlighting their complexity and paving the path towards dissecting multi-layered MTC functions in mammals.

## Introduction

The *N6*-methyl-adenosine (m6A) modification is the most widespread RNA modification on mammalian messenger RNA (mRNAs). The mark is deposited by a multi-subunit protein complex, also known as the m6A <u>m</u>ethyltransferase (“writer”) <u>c</u>omplex (MTC), of which the catalytic subunit is METTL3 in human and Ime4 in yeast, (Balacco & Soller, 2019; Clancy *et al*, 2002). In mammals, the deposition of m6A occurs at DRACH motifs throughout transcripts that are distant from splice sites, biasing this modification towards long internal and last exons (Dominissini *et al*, 2012; Meyer *et al*, 2012; Uzonyi *et al*, 2023). The presence of m6A on mRNA primarily impacts mRNA stability but has also been linked to a wide array of additional molecular outcomes, and its disruption is associated with diverse phenotypes ranging from immune dysfunction to cancer (Jiang *et al*, 2021; Murakami & Jaffrey, 2022; Zaccara *et al*, 2019).

METTL3, which forms a hetero dimer complex with METTL14, is the catalytic unit of the MTC (Liu *et al*, 2014; Wang *et al*, 2016). Several auxiliary proteins that interact with METTL3/METTL14 are required for the deposition of m6A. These include the scaffold proteins WTAP and VIRMA, E3 ligase HAKAI, ZCH13H3, and RNA binding proteins RBM15/RBM15b (Bawankar *et al*, 2021; Knuckles *et al*, 2018; Ping *et al*, 2014; Schwartz *et al*, 2014; Wang *et al*, 2021; Wen *et al*, 2018; Yue *et al*, 2018b). Structural and topological analyses of the mammalian and Drosophila MTCs revealed that MTCs can be divided into two subcomplexes: (1) the m6A METTL complex (MAC) which consists of METTL3/METTL14, and a (2) m6A-METTL associated complex (MACOM) consisting of WTAP, VIRMA, HAKAI, ZCH13H3, and RBM15/RBM15b (Figure 1A) (Lence *et al*, 2019; Su *et al*, 2022). MACOM directly interacts with MAC to enhance m6A deposition. In *Arabidopsis thaliana* all subunits of both MAC and MACOM, except for RBM15/RBM15b, are conserved and play related roles in m6A deposition suggesting structural and topological conservation in plants (Ruzicka *et al*, 2017; Zhang *et al*, 2022).

**Figure 1.**
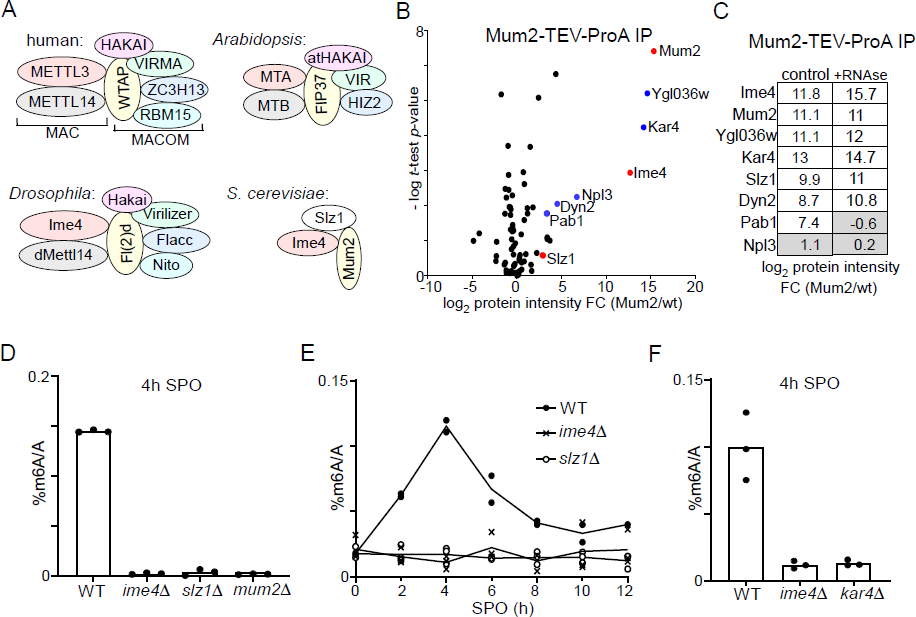
Identification of Mum2 interacting proteins. (**A**). Scheme of MTCs in mammals, Drosophila, Arabidopsis and yeast. Colour indicates matching orthologues. (**B**) Volcano plots of Mum2-TEV-ProA compared to untagged control. Diploid cells harbouring Mum2 tagged with TEV-ProA (FW7873) or untagged control (FW1511) were induced to enter meiosis. Protein extracts were incubated with ProA coated paramagnetic beads. TEV protease was used to elute Mum2 from the beads. Significantly enriched proteins are labelled in blue and known subunits of the MIS complex are labelled in red. (**C**) Similar analysis as in B, except that during IP extracts were either untreated (left panel) or RNase treated (right panel). (**D**) m6A levels as determined by LC-MS in WT diploid cells, or diploid cells harbouring *ime4*Δ, *mum*2Δ, or *slz1*Δ (FW1511, FW7030, FW6535, and FW6504). In short cells were grown to saturation in rich medium (YPD), shifted to BYTA and grown for another 16 hours, then shifted to sporulation medium (SPO). Samples were taken at 4 hours in SPO. Total RNA was extracted and followed by two rounds of polyA RNA purification. Subsequently, RNA was digested into nucleosides and m6A was quantified my LC-MS. The mean of n=3 biological replicates are shown. (**E**) Same analysis as in D, except that multiple time points after shifting cells to SPO were taken (0, 2, 4, 6, 8, 10, 12 hours in SPO). WT, *slz1*Δ or *ime4*Δ diploid cells were used. The mean values of n=2 biological repeats are shown. (**F**) m6A levels in WT, *ime4*Δ and *kar4*Δ cells (FW1511, FW7030, FW8246). Samples were collected at 4 hours in SPO. The mean of n=3 biological replicates are shown.

In yeast, m6A is deposited solely during early meiosis (Agarwala *et al*, 2012; Shah & Clancy, 1992). Intriguingly, early dissections of the yeast MTC have revealed only three proteins <u>M</u>um2, Ime4 and <u>S</u>lz1, collectively coined the MIS complex (Agarwala *et al*., 2012) (Figure 1A). However, only two of these proteins (Ime4 and Mum2) have mammalian homologs (METTL3 and WTAP respectively), which could suggest that MTCs have undergone substantial rewiring throughout evolution.

More than 1000 mRNAs are methylated during early yeast meiosis (Schwartz *et al*, 2013). Like in mammals, m6A modified mRNAs are turned over more rapidly compared to unmodified mRNAs (Bushkin *et al*, 2019; Scutenaire *et al*, 2022; Varier *et al*, 2022). The decay of m6A transcripts requires active translation and is mediated by the YTH domain containing protein Pho92/Mrb1 (Varier *et al*., 2022). Several studies have shown that preventing the m6A modification at single meiotic transcripts can cause defects in meiosis (Bushkin *et al*., 2019; Scutenaire *et al*., 2022). However, the exact function of the m6A modification in yeast meiosis remains elusive. In yeast, Ime4 has both m6A-dependent and m6A-independent functions in meiosis, as a catalytically inactive mutant of Ime4 shows milder delay in meiotic progression compared to the Ime4 deletion mutant (Agarwala *et al*., 2012; Clancy *et al*., 2002).

The fact that MTCs in mammals, Drosophila and plants harbour at least six conserved protein subunits (Figure 1A), motivated us to re-examine the composition of the yeast MTC. Here we uncover and characterize three novel components of the yeast MTC (Kar4, Ygl036w/Vir1 and Dyn2). We further demonstrate that Kar4 and Vir1, and potentially also Slz1, a previously known component, are orthologs of known components of the mammalian MTC (METTL14, VIRMA, and ZCH13H3, respectively). In total five members of the yeast MTC likely have orthologs matching the mammalian MTC. We show that Ime4, Mum2, Vir1 and Kar4 form a stable complex on mRNAs required for meiotic progression, while the Slz1 subunit is essential for m6A deposition but not for MTC integrity. The composition of the yeast MTC is considerably more similar to its mammalian counterparts than previously thought. Our findings also suggest that in contrast to mammals and Drosophila, the yeast MTC has no MAC and MACOM arrangement.

## Results

### Identification of Mum2 (and Kar4) interacting proteins

The inconsistency in the composition of the yeast MTC, comprising only three known components (Agarwala *et al*., 2012), in comparison to its mammalian, Drosophila and plant MTC counterparts all comprising at least six conserved proteins (seven in the case of mammals and Drosophila), prompted us to re-evaluate the yeast MTC using a proteomics approach (Figure 1A). We generated a yeast strain with the conserved scaffold protein in MTCs, Mum2, fused to the ProA-tag followed by a TEV cleavage sequence at the carboxy terminus. Subsequently, we performed immunoprecipitation with the ProA-tag in protein extracts generated from cells staged in early meiosis followed by elution with the TEV protease (Figures 1B, and Table S1). As expected, Ime4 was strongly enriched in the Mum2 IP-MS analysis. We also identified two other proteins that were strongly enriched, Ygl036w and Kar4. Two additional RNA binding proteins, Pab1 and Npl3, and light chain dynein protein, Dyn2, were significantly enriched in the Mum2 IP, albeit to a lesser extent (log_2_ protein intensity fold change (FC) >2, and −log *t*-test *p*-value >1). Noteworthy, Slz1, a known component of the MIS complex, was not significantly enriched (Figure 1B).

To confirm that Kar4 and Ygl036w are integral parts of the yeast MTC, we performed a Kar4 IP-MS using a similar setup (Figure S1A). Even though Kar4 enrichment was only 4-fold over background, we found that Mum2, Ime4 and Ygl036w, but not Slz1, significantly co-purified with Kar4 (log_2_ protein intensity FC >2, and −log *t*-test *p*-value >1).

Next, we examined whether the interaction with Mum2 was dependent on the presence of RNA in the sample by treating the protein lysate with RNase. The Mum2-IP MS analysis identified Ime4, Ygl036w, Kar4, Slz1, and Dyn2 in both control and RNAse-treated samples (Figures 1C, S1B, and Table S1). Pab1 was identified in the control but was not enriched in the RNAse treated sample, while Npl3 was not enriched at all in these analyses (log_2_ protein intensity FC >2, and −log *t*-test *p*-value >1) (Figures 1C, S1B, and Table S1). We conclude that Ime4, Ygl036w, Kar4, Slz1, and Dyn2 interact with Mum2 in an RNA independent manner, while Pab1 interaction with Mum2 is RNA-dependent and hence indirect. These data suggest that Kar4, Ygl036w and Dyn2 are MTC components.

### Kar4, Ygl036w and Slz1 are essential for m6A deposition

Next, we determined whether known members and newly identified interactors are required for m6A deposition. Previous work showed that m6A deposition is severely reduced in deletion mutants of two components of the MIS complex: *IME4* and *MUM2* (Agarwala *et al*., 2012). To confirm these previous findings, we determined m6A levels in diploid cells undergoing early meiosis (4 hours in sporulation medium (SPO)) harbouring gene deletion in *MUM2* and *IME4* (*mum2*Δ and *ime4*Δ). Both deletion mutants showed a severe reduction in m6A over A levels as determined by LC-MS (Figure 1D), consistent with previous measurements (Ensinck *et al*, 2023). We also quantified m6A levels in a *SLZ1* deletion strain. We found that *slz1*Δ cells showed a similar reduction in m6A levels as *ime4*Δ cells (LC-MS), suggesting that Slz1 is also required for m6A deposition (Figure 1D). The analysis contrasts previous work that showed only a partial reduction in m6A levels in *slz1*Δ cells (Agarwala *et al*., 2012). One possibility for the discrepancy is that *slz1*Δ cells had a delay in m6A accumulation during early meiosis, and possibly m6A accumulation occurs later in meiosis in *slz1*Δ cells. However, m6A levels in *slz1*Δ cells did not accumulate at any time in the 12 hours following meiotic induction and showed background levels similar to *ime4*Δ cells (Figure 1E).

In yeast, Kar4 is known to act as a transcription factor important for the mating response pathway, however no clear mechanistic role for Kar4 in meiosis has been reported (Kurihara *et al*, 1996). Interestingly, high-throughput studies suggest that the *KAR4* deletion negatively affects meiosis and sporulation (Deutschbauer *et al*, 2002; Enyenihi & Saunders, 2003). Based on sequence homology analysis, Kar4 has been proposed to be the yeast orthologue of METTL14, a critical component of MTCs in mammals, Drosophila, and plants. Phylogenetic analysis of methyltransferase domains showed that Kar4 and METTL14 belong to the same subfamily, but are relatively different from each other (Bujnicki *et al*, 2002). Since METTL14 is considered a structural subunit in the METTL3/METTL14 heterodimer, we hypothesized that Kar4 may play a related role in m6A deposition (Sledz & Jinek, 2016; Wang *et al*., 2016). We found that like *ime4*Δ cells entering meiosis, *kar4*Δ cells showed no detectable m6A levels (Figure 1F).

To examine whether Ygl036w, Dyn2 and the other identified Mum2 interacting proteins (Pab1 and Npl3) are important for m6A deposition, we performed m6A-ELISA and m6A-seq2 (Figure 2). Both analyses by m6A-ELISA and m6A-seq2 showed that *ygl036w*Δ had no detectable m6A levels, similar to *ime4*Δ (Figure 2A and 2B). Additionally, *dyn2*Δ had reduced m6A levels (50% of WT) as detected by m6A ELISA (Figure 2C). Pab1 is an essential gene and Npl3 is essential for entry into meiosis. To examine whether the two are required for m6A deposition, we depleted both proteins using the auxin induced degron (AID) system during meiosis (Figure S2). To obtain efficient depletion in meiosis, we let cells enter early meiosis (2 hours SPO) and subsequently induced depletion of Pab1 (*PAB1-AID*) or Npl3 (*NPL3-AID*) for 2 hours (Figure S2). As a control we also depleted Slz1 (*SLZ1-AID*). We found that depletion of Pab1 and Npl3 showed no significant reduction in m6A levels compared to the control (Figure 2D). As expected, depletion of Slz1 showed a strong reduction of m6A. Also given that Pab1 and Npl3 interactions with Mum2 were RNA dependent or not reproducible (in the case of Npl3), both RNA binding proteins are likely not part of the yeast MTC. We conclude that in addition to Ime4 and Mum2, Slz1, Kar4, and Ygl036w are all essential for m6A deposition.

**Figure 2.**
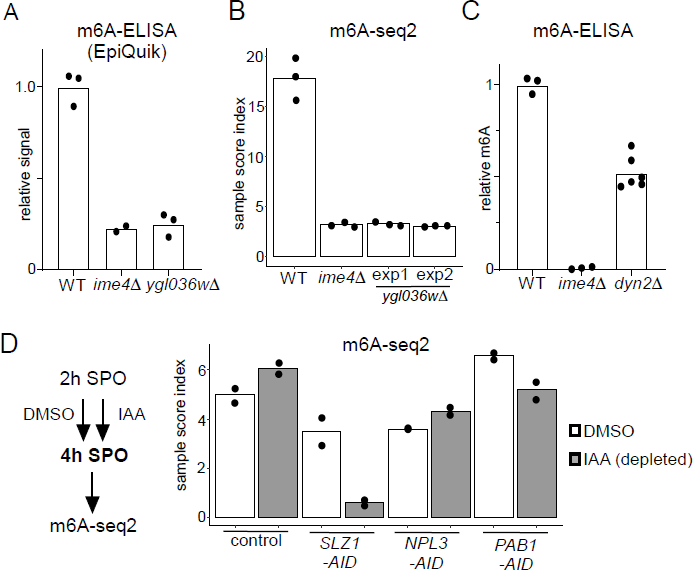
Functional characterization of Mum2 interactors. (**A**) WT, *ime4*Δ, and *ygl036w*Δ cells (FW1511, FW7030, FW9307) were induced to enter meiosis. RNA was extracted, and polyA RNA was purified. m6A levels were determined by m6A-ELISA kit supplied by Epiquik. The means of n=3 biological replicates are shown. (**B**) m6A-seq2 analysis of strains and condition described in A. Two independent clones of *ygl036w*Δ were used for the analysis. Shown are the sample index score representing 1308 known of m6A sites. The mean of n=3 biological repeats are shown. (**C**) m6A levels WT, *ime4*Δ, and *dyn2*Δ cells (FW1511, FW7030, FW10442). RNA was extracted, and polyA RNA was purified. m6A levels were determined by a non-commercial m6A-ELISA assay (Ensinck *et al*., 2023). The means of n=3 biological replicates are shown of WT and *ime4*Δ, and n=6 for *dyn2*Δ. (**D**) m6A-seq2 analysis after Npl3, Pab1 and Slz1 depletion. Diploid cells containing *PAB1-AID*, *NPL3-AID*, *SLZ1-AID* (FW10388, FW10389, FW10386) were treated at 2 hours in SPO for 2 hours with DMSO or IAA and CuSO4 for rapid depletion. A control strain was included that only harboured the TIR1 ligase expressed from the *CUP1* promoter (FW5737). Shown are the m6A-sample indices, quantifying the overall levels of enrichment over 1308 previously defined m6A sites. The mean of n=2 biological repeats are shown.

### The m6A writer complex is conserved

Our analysis revealed that the yeast MTC comprises at least six protein subunits (Mum2, Ime4, Kar4, Ygl036w, Slz1, Dyn2), all of which - with the exception of Dyn2 - are required for m6A methylation. Two subunits that were described previously have mammalian orthologues: Ime4/METTL3 and Mum2/WTAP. Among the newly identified subunits Kar4 has a clear mammalian orthologue, named METTL14.

The two remaining subunits, Ylg036w and Slz1, did not have any obvious homologs based on standard BLAST searches. To investigate this further, we first conducted pairwise comparisons of the Ylg036w protein coding sequence against all sequences of the human proteome, relying on both global (Needle) and local (Matcher) sequence similarity scores (Figure 3A). The analysis revealed that Ygl036w displayed a striking sequence similarity with VIRMA, a known subunit of the mammalian MTC. We next performed proteome-wide structural alignments of the predicted structure of Ygl036w against all predicted structures of the human proteome. The top hit in this analysis, by far, was VIRMA (Figure 3B). A structural alignment of Ygl036w against VIRMA displays a high extent of agreement (TM-score of 0.483, normalized for Ygl036w) (Figure 3C and S3). These analyses thus strongly suggest that Ygl036w and VIRMA are homologs, and we hence renamed Ygl036W to Vir1, as the yeast orthologue of VIRMA in mammals, Virilizer in Drosophila, and VIR in plants, respectively.

**Figure 3.**
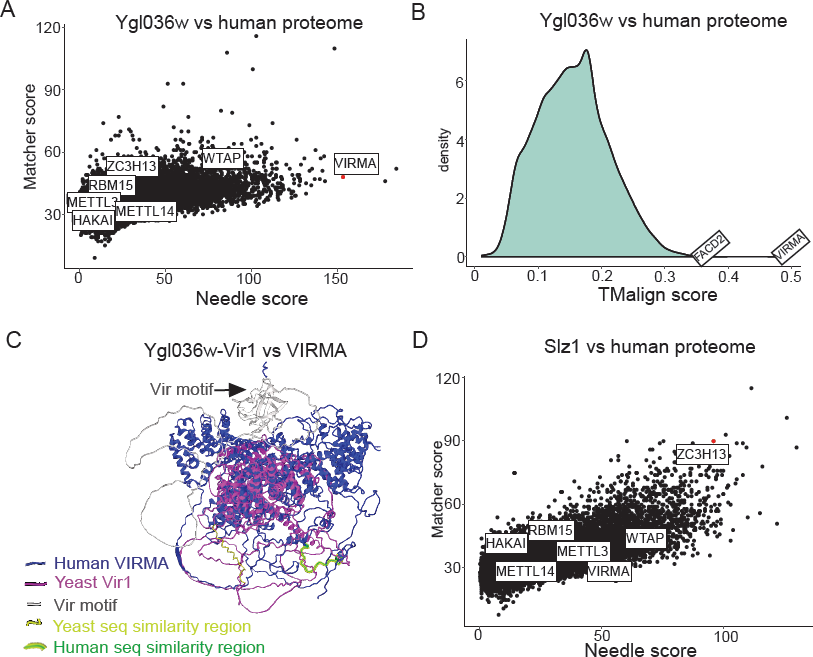
Ylg036w-Vir1 and Slz1 have orthologues in humans. **(A)** Pair-wise sequence comparison of Ylg036w against all human protein coding genes. Global similarity scores (using Needle) are displayed on the x-axis, whereas local similarity scores (using Matcher) are displayed on the y-axis. **(B)** Pairwise structural similarity of Ygl036w against all human protein coding genes, using TMalign algorithm. A histogram of the TMalign density scores is displayed, along with an indication of the score against VIRMA, the top hit in this analysis, and the second highest hit, FACD2. TM-scores were calculated and normalized using the query gene length, Ygl036w. **(C)** Overlay of Alpha-fold predicted structures of Ylg036c-Vir1 and human VIRMA. Human VIRMA is shown in blue and Yeast Ylg036c-Vir1 is shown in pink. Also indicated are the Vir motif and regions with strong sequence similarities. **(D)** Analysis as in A using Slz1 as a query.

Motivated by these results, we next sought to conduct the same sets of analyses for Slz1. Strikingly, Slz1 showed partial similarity with ZC3H13 (Figure 3D). ZC3H13 is highly unstructured, and much larger than Slz1, which made 3D structural alignments uninformative. Interestingly, both SLZ1 and ZC3H13 have been proposed to play a similar role, of shuttling the MTC complex into the nucleus (Schwartz *et al*., 2013; Wen *et al*., 2018), providing further evidence that these two proteins may be orthologues. We conclude that the yeast MTC is considerably more conserved than previously described, with five subunits of the mammalian MTC complex (METTL3, METTL14, WTAP, VIRMA, ZC3H13) likely having orthologues in the yeast counterpart (Ime4, Kar4, Mum2, Vir1, Slz1).

### Topological features of the yeast MTC

To gain further insight on the organization of the yeast MTC, we dissected the interrelationships between the MTC components. Often, protein subunits interacting as stable macromolecular complexes are destabilized when protein complexes are disrupted, e.g. upon deletion of a scaffold protein. With this in mind, we determined protein expression levels of all yeast MTC subunits in deletion mutants for each of the remaining subunits (Figures 4A and S4A-E). The analyses revealed that Ime4, Mum2 and Vir1 were all required for mutual stabilisation in cells undergoing meiosis, with protein levels of any of these three components being strongly reduced upon elimination of any of the other two (Figures 4A, 4B, and S4A-C). Interestingly, loss of any of these proteins also led to loss of Kar4, yet - surprisingly - this was not mutual, as *kar4*Δ had little effect on expression of any of the subunits (Figures 4A, 4B, and S4D). The *slz1*Δ had little effect on expression of any of the subunits (Figures 4A, 4B, and S4A-D). Importantly, loss of the different components was not due to changes in RNA levels as shown by RNA-seq, as these remain relatively unchanged in the mutants except for Slz1, where *SLZ1* mRNA induction was reduced in the deletion mutants resulting in lower Slz1 protein levels (Figures 4A, 4B, S4E and S4F). These results are at odds with results in mammalian cells where METTL14, the Kar4 orthologue, stabilises METTL3, and VIRMA is not important for METTL3 and METTL14 stabilization (Wang *et al*., 2016; Wang *et al*, 2014; Yue *et al*., 2018b).

**Figure 4.**
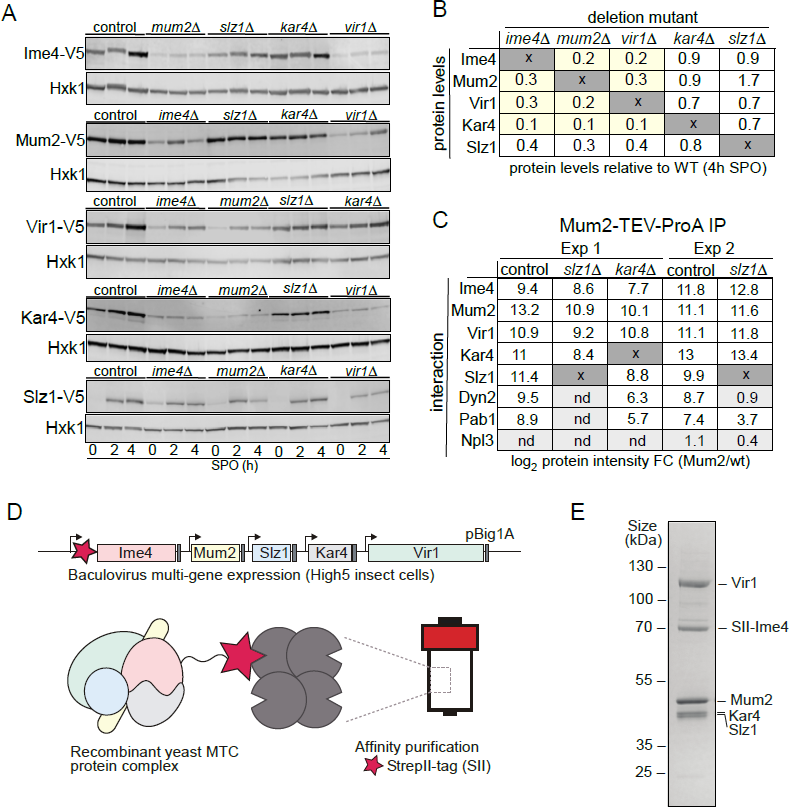
Topological and structural properties of the yeast MT. **(A)** Ime1, Mum2, Vir1, Kar4, and Slz1 expression depends on the presence of MTC components. WT, *mum2*Δ, *ime4*Δ*, slz1*Δ*, kar4*Δ or *vir1*Δ were examined for Ime4-V5 (first panel, FW6057, FW8362, FW8264, FW8633, and FW9483), Mum2-V5 (second panel, FW6500, FW9394, FW6534, FW9396, and FW9398), Vir1-V5 (third panel, FW9666, FW9663, FW9668, FW9670, and FW9643), Kar4-V5 (fourth panel, FW8216, FW9484, FW8212, FW9482, and FW9481), or Slz1-V5 (fifth panel, FW6502, FW9479, FW9477, FW9480, and FW9478) during early meiosis. Samples were taken at the indicated time points. Western blots were probed with anti-V5 antibodies and anti-Hxk1 as a loading control. (**B**) Table of western blot quantifications of the 4-hour time point described in C. Each row represents the protein expression (Ime4-V5, Mum2-V5, Vir1-V5, Kar4-V5 and Slz1-V5) and each column the deletion mutant (*mum2*Δ, *ime4*Δ*, slz1*Δ*, kar4*Δ or *vir1*Δ). Highlighted in yellow are the MTC components affected in protein levels (<0.5), but not at the mRNA levels. (**C**) Diploid cells untagged or harbouring Mum2 tagged with TEV-ProA in WT, *slz1*Δ, or *kar4*Δ background (FW1511, FW7873, FW10158, and FW10159) were induced to enter meiosis. Protein extracts were incubated with ProA coated paramagnetic beads. TEV protease was used to elute Mum2 from the beads. Shown are the enrichments for Ime4, Mum2, Vir1, Kar4, Slz1, Pab1, and Npl3. Two independent experiments are shown for *slz1*Δ. nd= not detected. **(D)** Purification of the 5-subunit recombinant m6A writer complex. Schematic representation showing the baculovirus recombinant co-expression and purification of the yeast MTC with five subunits (Ime4, Mum2, Vir1, Kar4, and Slz1) in insect cells. Genes are depicted by rectangles, promoters by arrows and terminators by grey boxes. The star in magenta represents the Strep-tag in the N-terminal domain of Ime4. **(E)** SDS-PAGE analysis of the purified recombinant m6A writer complex after affinity. Identities of bands confirmed by MS are labelled.

To determine the manifestation of these interdependencies in the composition of the m6A writer complex, we performed Mum2 IP-MS in WT, *slz1*Δ and *kar4*Δ cells (Figures 4C, and Table S1). In line with the protein expression analysis of the m6A writer complex deletion mutants, we found that in *slz1*Δ cells the Mum2-Ime4-Vir1-Kar4 complex remained intact, and in *kar4*Δ cells Mum2 formed interactions with Vir1 and Ime4. Slz1 and Dyn2 were also both enriched in the Mum2-IP in *kar4*Δ cells, suggesting that Kar4 does not mediate interactions between the other subunits of the m6A writer complex (Figure 4C and Table S1). Interestingly, Dyn2 was not identified as an interactor with Mum2 in the *slz1*Δ cells, suggesting that Slz1 mediates the interaction of Dyn2 with the MTC (Figure 4C and Table S1).

Our analyses showed that in cells undergoing meiosis Ime4, Mum2, Vir1, Kar4, Slz1 and Dyn2 make up the MTC, and that, except for Dyn2, all MTC components are essential for m6A deposition. However, it does not exclude the possibility that other proteins might be required for MTC assembly. Therefore, we co-expressed all five subunits essential for m6A (Ime4, Kar4, Mum2, Vir1, Slz1) in insect cells. Subsequently, we performed a single step affinity purification using the Strep-tag system with a Strep-tag II fused to the amino-terminal domain of Ime4 (Figure 4D). The analyses revealed that Vir1, Mum2, Kar4, and Slz1 co-purified with Ime4 (Figure 4E), showing that these five subunits form a stable complex *in vitro*. Collectively, these analyses suggest that the yeast MTC comprises three mutually stabilizing components: (1) a core comprising Ime4, Mum2, and Vir1, (2) Kar4, which also is stabilized when part of the MTC, and (3) a module comprising Slz1 and Dyn2, where Slz1 acts as a hub for binding of Dyn2. Additionally, we showed that the MTC components, Ime4, Mum2, Vir1, Kar4 and Slz1, form a stable complex *in vitro*.

### The yeast MTC has m6A dependent and independent functions in meiosis

It has previously been established that Ime4, the catalytic component, has both m6A dependent and m6A independent functions, on the basis of the observation that an Ime4 catalytic inactive mutant (*IME4*^CD^) displays a milder phenotype than an *IME4* deletion mutant (Agarwala *et al*., 2012; Clancy *et al*., 2002). We confirmed the difference between *ime4*Δ and *IME4*^CD^. Indeed, we found *IME4*^CD^ showed no m6A deposition and a milder delay in meiosis compared the *ime4Δ* (Figure S5A and S5B). One can conceive of two classes of m6A independent roles for Ime4: (1) A ‘moonlighting’ function, independent also of the role of Ime4 in forming part of the yeast MTC, (2) an MTC-dependent (yet m6A-independent) function. We reasoned that the availability of three (Vir1, Kar4, Dyn2) new additional components of the MTC complex might allow us to distinguish between these possibilities. If the m6A independent role of Ime4 is independent of the MTC, deletion of *IME4* would result in a phenotype unique to Ime4 and not overlapping with its counterparts upon deletion of other MTC components. In contrast, if the phenotype is due to the requirement of the MTC complex, identical phenotypes should be observed also upon loss of additional components.

Accordingly, we monitored the progression of meiosis upon loss of each of the MTC components. *kar4*Δ, *vir1*Δ, *mum2*Δ, and *ime4*Δ cells all displayed a strong defect in meiosis, and less than 40 percent completed meiotic divisions after 24h in SPO (Figure 5A). In contrast, *slz1*Δ had a considerably milder phenotype in meiosis compared to *ime4*Δ cells (Figure 5A) (Agarwala *et al*., 2012). *dyn2*Δ showed the mildest phenotype in meiosis with only marginal delay in meiotic divisions and little effect on the number of cells that underwent meiosis after 24h in SPO (Figure 5B).

**Figure 5.**
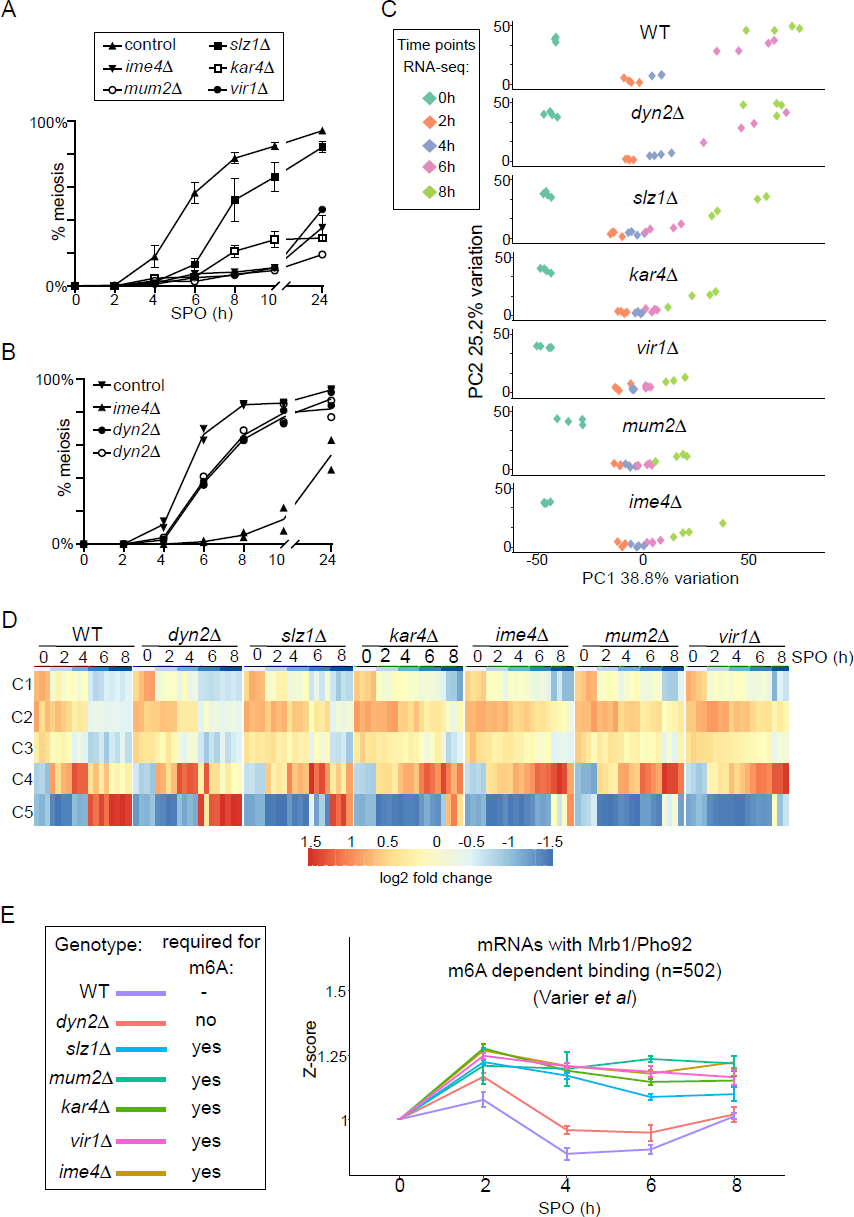
m6A dependent and independent roles of the MTC in meiosis. **(A)** Onset of meiosis in WT, *ime4*Δ, *mum2*Δ, *slz1*Δ, *kar4*Δ, and *vir1*Δ (FW1511, FW7030, FW6535, FW6504, FW8246, and FW9307). Cells induced to enter meiosis. Samples were taken at the indicated time points for DAPI staining. Cells were fixed, stained with DAPI, and nulei were counted for at least 200 cells per biological repeat. Cells with two or more DAPI masses were considered to undergo meiosis. The mean and SEM of n=3 biological repeats are displayed. **(B)** Similar analysis as A, except that the WT, *ime4*Δ, and *dyn2*Δ were analysed (FW1511, FW7030, FW10442 and FW10443). For the analysis two independent *dyn2*Δ strains were used. The mean of n=2 biological repeats are displayed. **(C)** Principal component analysis (PCA) of gene expression measurements across diverse strains and timepoints, done using PCAtools 2.6.0 for R. Strains described in A and B were induced to enter synchronized meiosis and samples were taken at the indicated time points (0, 2, 4, 6, and 8 hours in SPO). For analysis n=4 biological repeats were used, except for *ime4*Δ_4h, *ime4*Δ_0h, and *vir1*Δ_8h which had an n = 3. **(D)** Median expression patterns in WT and MTC mutants of five predefined gene clusters representing early metabolic genes (C1, n=67), early meiotic genes (C2, n=42; C3, n=34), early to middle meiotic genes (C4, n=53), middle meiotic genes (C5, n=136) (Chu *et al*., 1998). **(E)** Median gene expression fold change from timepoint 0 of Mrb1/Pho92-bound transcripts during the time course across the different deletion mutants. Dataset of Mrb1/Pho92 iCLIP-based targets associating with RNA in an m6A-dependent manner identified in (Varier *et al*., 2022) was used for the analysis. For each transcript a Z-score was calculated with respected to the 0-hour timepoint. The mean of at least n=3 biological replicates shown, and error bars represent the standard error of the mean.

To characterize the role played by the different writer complex subunits during meiosis in greater detail, we profiled RNA expression in each of the MTC mutants along a high-resolution meiotic time course. We took samples at 0, 2, 4, 6 and 8 hours following meiotic induction, which covers the early and middle phases of the yeast sporulation programme. For the analysis, we compared WT, *ime4*Δ, *mum2*Δ, *slz1*Δ, *kar4*Δ, *vir1*Δ and *dyn2*Δ. The experiment allowed us to assess more precisely which stage of meiosis was affected in the different deletion mutants of the m6A writer complex (Table S2). Two approaches were taken to analyse the data. First, we used a principal component analysis (PCA) and hierarchical clustering across all time points and all mutants (Figure 5C and S5C). Second, we assessed gene expression across different gene clusters in meiosis described previously (Chu *et al*, 1998). These genes clusters represent the early metabolic genes (cluster C1), early meiotic genes (cluster C2 and C3), early to middle meiotic genes (cluster C4), and middle meiotic genes (cluster C5), respectively (Figure 5D). Consistent with the meiotic phenotypes, we found that *kar4*Δ, *vir1*Δ, *mum2*Δ, and *ime4*Δ displayed the most severe expression phenotypes (Figures 5C, 5D, and S5C). This is most pronounced between the 4-to-6-hour timepoints, where *kar4*Δ, *vir1*Δ, *mum2*Δ, and *ime4*Δ remained staggered at cluster C2, whereas WT transitioned into cluster C4 and C5. The fact that the different MTC mutants clustered together is highly suggestive of a function common to all of them, as opposed to moonlighting functions of individual components (Figure 5C, 5D, and S5A). Also consistent with the phenotypic analysis, *slz1*Δ showed a mild delay, distinct from the remaining mutants, whereas *dyn2*Δ showed little difference in comparison to WT (Figures 5C, 5D and S5C).

Interestingly, despite the general lack of differences between the strains at the 2-hour timepoint, the histone genes, *HHF2* and *HTB2*, were significantly reduced in expression in five of the deletion mutants (*ime4*Δ, *mum2*Δ, *slz1*Δ, *kar4*Δ, *vir1*Δ), potentially hinting at an early role of the MTC complex in regulation of histone levels, reminiscent of recent findings in planaria (Figure S5D) (Dagan *et al*, 2022).

Finally, we determined whether the m6A-dependent decay function is reflected in the RNA-seq data set. Recent work showed that the sole m6A reader protein in yeast Mrb1/Pho92 controls the decay of m6A modified mRNAs during meiosis (Varier *et al*., 2022). We used an iCLIP dataset that identified the mRNAs associated by Pho92 in an m6A dependent manner to assess how these mRNAs were affected throughout the time course in WT and deletion mutants (Varier *et al*., 2022). Using this subset of the transcripts, we observed that in WT cells, these transcripts were generally induced up to the 2-hour timepoint, following which their levels declined. In contrast, in all five deletion mutants essential for m6A (*ime4*Δ, *mum2*Δ, *slz1*Δ, *kar4*Δ, *vir1*Δ), their levels failed to decline after their initial induction at 2 hours (Figure 5E). Consistent with the m6A and meiotic phenotype, the *dyn2*Δ only showed marginal difference compared to the WT (Figure 5E). A control set of randomly chosen transcripts did not display a difference between the WT and the deletion mutants (Figure S5E). These data support the m6A-dependent function of the MTC complex subunits, which is to m6A modify mRNAs for decay via Pho92 during early yeast meiosis (Varier *et al*., 2022).

Taken together, these analyses suggest that Kar4, Mum2, Vir1, and Ime4 play a dual role in controlling the expression program of meiosis via both m6A-dependent and m6A-independent yet MTC-complex-dependent functions, in contrast to Slz1 exhibiting only an m6A dependent function.

### The molecular functions of Kar4 in meiosis and mating differ

Kar4 has been shown to be important for controlling transcription of genes involved in the mating pathway in yeast. When haploid cells sense mating pheromone, Kar4 is induced and is recruited to promoters predominantly controlled by the transcription factor Ste12 to activate transcription of genes involved in the mating pathway (Gammie *et al*, 1999; Kurihara *et al*., 1996; Lahav *et al*, 2007). Our above-described findings reveal that Kar4 is also an integral component of the yeast MTC and important for meiosis. We therefore wondered whether the mating and meiotic functions might be mediated via a shared mechanism. Under such a scenario, Kar4’s roles in mating might require the additional MTC components, and - conversely – the role of Kar4 in meiosis might be mediated via binding to chromatin. Under the latter scenario, we speculated that Kar4 might function in directing the MTC to chromatin to facilitate co-transcriptional m6A deposition, which could be consistent with one of the proposed modes of action of METTL14 in mammals (Huang *et al*, 2019). To examine this, we first tested whether Ime4, Slz1, or Mum2 are expressed during mating. As expected, a shorter protein isoform of Kar4 was strongly induced when *MAT*a cells were treated with mating pheromone (α-factor) (Figure 6A) (Gammie *et al*., 1999). However, Ime4 and Mum2 expression were not elevated and Slz1 expression was not detected at all upon pheromone treatment. We also examined whether Kar4 expression during mating was dependent on the components of the MIS complex as observed in meiosis. We found that Kar4 levels were unaffected in *ime4*Δ, and *mum2*Δ cells (Figure 6B). In line with protein expression data, we found that Kar4 was bound to the *AGA1* promoter upon mating pheromone treatment as determined by ChIP, while other MTC subunits (Ime4, Mum2, Slz1) showed no enrichment (Figure 6C) (Aymoz *et al*, 2018). These data indicate that the yeast MTC does not play a role in the mating pathway.

**Figure 6.**
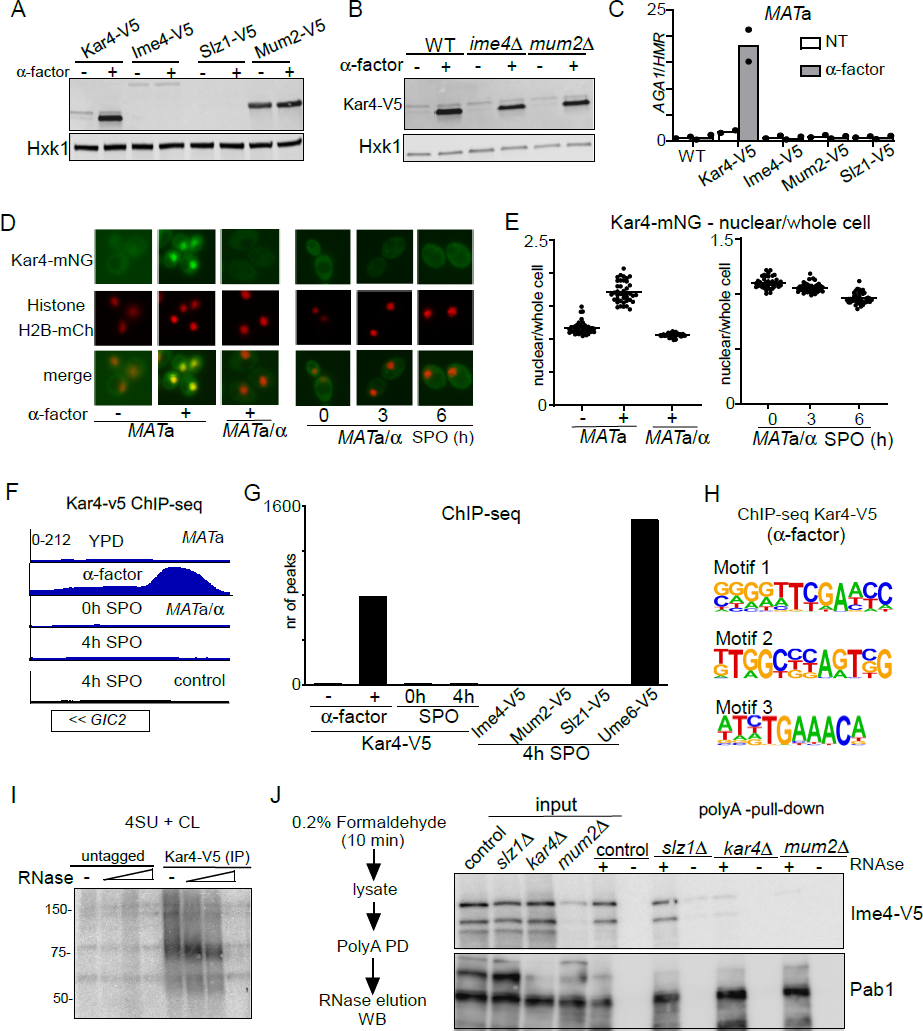
Kar4 functions during mating and meiosis differ. **(A)** Expression of Kar4, Ime4, Slz1, and Mum2, either untreated or treated with mating pheromone. *MAT*a harbouring V5 tagged alleles of Kar4, Slz1, Ime4, or Mum2 (FW8199, FW6426, FW5898 and FW6428) cells were grown in rich medium (YPD) until exponential growth, and then treated with α-factor for 30 minutes. Samples were collected for western blot and probed with anti-V5 antibodies. Hxk1 was used as a loading control. (**B**) Kar4 expression in WT, *ime4*Δ or *mum2*Δ *MAT*a cells (FW8199, FW9415, FW8574) untreated or treated α-factor. Samples were collected for western blot and probed with anti-V5 antibodies. Hxk1 was used as a loading control. (**C**) Chromatin immunoprecipitation (ChIP) of untagged control, Kar4-V5, Ime4-V5, Mum2-V5, and Slz1-V5 *MAT*a cells (FW1509, FW8199, FW5898, FW6426, and FW6428). Cells were grown in YPD and were either not treated or treated with α factor for 2 hours. (**D**) Kar4 localization in cells in rich medium, either untreated or treated α-factor and in cells entering meiosis. Cells were induced to enter meiosis in SPO, and samples were taken at 0, 3, and 6 hours in SPO. Kar4 fused to mNeongreen (mNG) was used for the analysis. To determine nuclear Kar4-mNG signal, we used histone H2B fused to mCherry (H2B-mCh). For the analysis we used *MAT*a and *MAT*a/α cells (FW8615 and FW8646). Representative images are shown. (**E**) Quantification of nuclear over whole cell mean signal for Kar4-mNG of data in C. At least n=50 cells were quantified for the analysis. (**F**) Chromatin immunoprecipitation followed by deep-sequencing (ChIP-seq) of *MAT*a Kar4-V5 (FW8199) untreated or treated with α factor, and *MAT*a/α Kar4-V5 (FW8616) at 0 and 4 hours in SPO as well as untagged control cells (FW1509 and FW1511). Samples were crosslinked with formaldehyde, extracts were sonicated, and protein-DNA complexes were purified using anti-V5 agarose beads and reverse crosslinked. Purified DNA was subjected to deep sequencing. ChIP-seq signals for the *GIC2* and *FUS1* loci are shown. (**G**) Quantification of the number of ChIP-seq peaks. (**H**) Motif analysis of Kar4 binding sites identified in Kar4-V5 ChIP-seq of α factor treated cells. **(I)** Photo activatable crosslinking of Kar4. Control and Kar4 (FW1511 and FW8633) cell entering meiosis were incubated with 4thiouracil (4TU). Cells were UV-crosslinked, and Kar4 was immunoprecipitated from protein extracts. The Kar4-IP was treated with different concentration of RNase, and subsequently radioactively labelled p32. Kar4-RNA complexes were separated by SDS page. **(J)** Chemical RNA-protein interactome analysis for assessing Ime4 binding to RNA. For the analysis we used Ime4-V5 cells in control, *slz1*Δ*, kar4*Δ*, mum2*Δ backgrounds (FW6057, FW8362, FW8264, and FW8633). In short, cells entering meiosis (4h SPO) were crosslinked with formaldehyde, polyA mRNAs were pulled down from protein extracts with oligo-dT coated magnetic beads, washed, samples were eluted with RNase, and assessed by western blotting. Membranes were probed for anti-V5 to detect Ime4. As a positive control, membranes were probed with anti-Pab1.

Second, we determined Kar4 localization during meiosis and mating by using a strain with Kar4 fused to mNeongreen (Kar4-mNG). We found that the Kar4 localization pattern was different between mating and meiosis. Kar4-mNG concentrated inside the nucleus in *MAT*a cells treated with α-factor (Figures 6D and 6E, and S6A). During meiosis (*MAT*a/α cells), however, we found that Kar4-mNG was localized to both the cytoplasm and the nucleus (Figures 6D, 6E, and S6B).

Third, we examined whether Kar4 or several components of the yeast MTC (Ime4, Mum2, and Szl1) associate with chromatin during meiosis (Figures 6F, 6G, S6C and S6D). While Kar4 was bound to 800 genomic locations in *MAT*a cells treated with α-factor, only few Kar4 peaks were detected during meiosis in *MAT*a/α cells (less than 10) (Figure 6F and 6G). As expected, Kar4 ChIP-seq peaks were enriched for the Ste12 binding sites (motif 3 in Figure 6H), but other motifs were also identified, perhaps indicating that Kar4 can bind to promoters in a Ste12 independent manner (Figures 6H and S6D). In line with Kar4, ChIP-seq of Ime4, Mum2, and Slz1 showed no enrichment throughout the genome during meiosis (Figures 6G and S6C). As a control we included ChIP-seq of the key meiotic transcriptional regulator Ume6 under the same conditions, which showed, as expected, enrichment at over 1000 genomic locations (Figure 6G) (Chia *et al*, 2021). Lastly, we examined whether the Ste12 transcription factor, which is important for Kar4 recruitment during mating, plays a role in meiosis (Kurihara *et al*., 1996; Lahav *et al*., 2007). We found that *ste12*Δ did not affect the onset of meiosis (Figure S6E). The above results thus rule out the requirement of additional MTC components during mating, and further rule out a Kar4-mediated association of the yeast MTC with chromatin during early meiosis. We conclude that the Kar4 function in the MTC during meiosis differs from its known role as transcription regulator in the mating response pathway.

### Kar4 stabilizes the MTC on mRNAs

Given the different roles of Kar4 in meiosis and mating, and that Kar4 was not required for stabilization of Ime4, we sought to better understand the molecular role played by Kar4 in the MTC complex during meiosis. Specifically, we examined whether Kar4 was required for allowing Ime4 to associate with mRNAs.

We assessed RNA binding of the m6A writer complex in cells staged in early meiosis. We applied RNA-protein interaction capture after chemical crosslinking to purify protein-RNA complexes from cells after short and mild crosslinking with formaldehyde using oligo(dT) magnetic beads (Figure S6F). The assay revealed that both Ime4 and Kar4 efficiently co-purify with mRNA following RNAse elution from the beads, while pulldown from extracts pre-treated with RNAse did not show Ime4 and Kar4 signals with immunoblot as expected (Figure S6F). As an orthogonal approach, we performed photo activatable UV crosslinking to determine whether RNA associate with Kar4. The analysis revealed Kar4 can crosslink to RNA (Figure 6I and S6G). Thus, Ime4 and Kar4 can both associate with mRNAs during early meiosis.

Next, we assessed whether Ime4 association with mRNAs is regulated by Kar4, Slz1, and Mum2 (Figure 6J). We applied RNA-protein interaction capture on extracts from *slz1*Δ, *kar4*Δ and *mum2*Δ cells that were induced to enter meiosis. After the polyA pull-down, and elution with RNase, Ime4 was specifically enriched to comparable degrees in both the control and *slz1*Δ (Figure 6J). In contrast, Ime4 association to mRNAs was reduced in *kar4*Δ cells with the no effect on Ime4 levels in input. Given Mum2’s stabilizing role in the yeast MTC, the Ime4 signal in *mum2*Δ cells was greatly reduced in both input and eluate from polyA pull-down. The Pab1 protein signal, which served as a positive control for RNA binding, did not differ across the different samples. These data suggest that Kar4 is important for stabilizing the interaction between Ime4 (and likely the whole MTC) and mRNAs, and that Slz1 is not required for this.

## Discussion

Previous work showed that the yeast MTC consists of only three subunits (Mum2, Ime4, Slz1), which contrasted the six or seven subunits described for MTCs in mammals, Drosophila, and plants, respectively (Agarwala *et al*., 2012). Here, we showed that the yeast MTC comprises six subunits of which five are essential for m6A deposition and have orthologues in the mammalian MTC. While the yeast MTC subunits are considerably more conserved than previously described, our findings suggest that the MTC is configured differently compared to the mammalian and Drosophila counterparts (see below). The yeast MTC exerts both m6A dependent and independent functions that are both crucial for yeast meiosis.

### Composition and features of the yeast MTC

Our analysis suggests that five subunits of the yeast MTC have orthologues in mammalian MTCs (Ime4/METTL3, Kar4/METTL14, Mum2/WTAP, Vir1/VIRMA, and potentially also Slz1/ZC3H13) (Figure 7). Thus, the composition of the yeast MTC is more conserved than previously described, however, we propose that the yeast MTC subunits are configured differently in a complex. The mammalian and Drosophila MTCs are configured into MAC (METTL3 and METTL14) and MACOM (WTAP, VIRMA, ZC3H13, HAKAI and RBM15) complexes, of which MAC harbours the catalytic activity and MACOM is critical for stimulating MAC activity (Knuckles *et al*., 2018; Su *et al*., 2022). Consistent with the presence of two complexes, diverse studies over the years have reported that METTL3 and METTL14 mutually stabilize each other (Sledz & Jinek, 2016; Wang *et al*., 2016). Likewise, subunits of the MACOM complex (e.g. WTAP and VIRMA) mutually stabilize each other (Yue *et al*, 2018a). However, depletion of components in one of subcomplex was not reported to impact the integrity of the other subcomplex (Yue *et al*., 2018a). In yeast, we show that five subunits (Mum2, Ime4, Vir1, Kar4, and Slz1) are essential for m6A deposition and form a stable complex. We also provide evidence that the yeast MTC subunits Ime4, Mum2, and Vir1 stabilize each other and Kar4, indicating that in yeast the MAC and MACOM subunits are interweaved (Figure 7). This interweaved arrangement of the yeast MTC is accompanied also by loss of a mutual stabilizing interaction between Ime4 and Kar4, which is in contrast to their mammalian counterparts (Sledz & Jinek, 2016; Wang *et al*., 2016).

**Figure 7.**
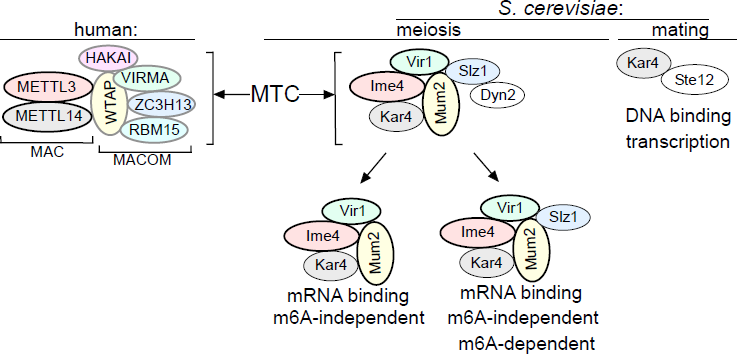
Model of the yeast MTC. Comparison of the yeast and the human MTCs. The human MTC consists of MAC and MACOM subcomplexes, while the yeast MTC forms a single complex. The conserved subunits are colour indicated. Mum2, Vir1, and Ime4 mutually stabilize each other. Further indicated are the m6A-dependent and m6A-independent MTC requirements, and Kar4’s separate function in mating.

We showed that Ygl036w-Vir1 is the orthologue of Virma in mammals, Virilizer in Drosophila, and Vir in plants, respectively (Haussmann *et al*, 2016; Ruzicka *et al*., 2017; Yue *et al*., 2018a). Like its mammalian/Drosophila/plant counterparts, Vir1 has a central role in the MTC and m6A deposition. Structurally, Vir1 and VIRMA showed strong conservation, however Vir1 does not contain the VIRMA domain which is thought to bridge the interaction between HAKAI and Fl2(d) in Drosophila (Bawankar *et al*., 2021; Wang *et al*., 2021). The absence of this domain may underlie the lack of a discernible HAKAI orthologue in yeast. We showed that in yeast, Vir1 stabilizes Mum2, Ime4, and Kar4, and that it is stabilized by Mum2 and Ime4. This contrasts with the mammalian MTC, where VIRMA and WTAP stabilize each other, but not METTL3 and METTL14 (Yue *et al*., 2018a).

Though Kar4 has been proposed to be the METTL14 orthologue in yeast, no molecular function in meiosis and in m6A deposition for yeast Kar4 was described previously. Here we showed that Kar4 is a critical subunit of the yeast MTC. Like METTL14, Kar4 is essential for m6A deposition and facilitates mRNA binding of the yeast MTC *in vivo*. We speculate that that the role of Kar4 in bridging mRNA binding of the yeast MTC is critical since in yeast the MTC likely does not harbour a designed RNA binding protein such as RBM15 (Patil *et al*, 2016). This function of Kar4 is perhaps related to how METTL14 stabilizes METTL3 on mRNAs *in vitro* and *in vivo* (Sledz & Jinek, 2016; Wang *et al*., 2016).

We found that Slz1 shows some sequence similarities with ZC3H13 in humans, suggesting Slz1 and ZC3H13 are possible orthologues despite being very different in size. Previous work showed that Slz1 is required for localizing the yeast MTC to the nucleus, which is also consistent with findings for ZC3H13 in mammals where it also is important for MTC nuclear localization (Schwartz *et al*., 2013; Wen *et al*., 2018). In Drosophila Flacc (ZC3H13/Slz1) bridges the interaction between Nito (RBM15) and the MTC (Knuckles *et al*., 2018). However, our analysis failed to identify a yeast orthologue for Nito (RBM15) (Knuckles *et al*., 2018). Slz1 does not seem to be important for MTC association with mRNAs and is dispensable for the m6A-independent role of the MTC. However, Slz1 is essential for driving the m6A deposition reaction since cells lacking Slz1 showed no m6A. Recent structural work on the human MTC showed that ZC3H13 facilitates a conformational change in VIRMA and WTAP, which may be important for m6A deposition (Su *et al*., 2022). The relevance of such a role in yeast remains to be confirmed.

Our analysis showed that dynein light chain protein, Dyn2, is a subunit of the yeast MTC. Despite the small size of Dyn2, we identified Dyn2 in all Mum2-IP-MS experiments. Moreover, the interaction between Dyn2 and the yeast MTC Mum2 is RNA independent but requires Slz1. In contrast to the remaining components, loss of Dyn2 led to a partial loss of mRNA methylation accompanied by a very mild meiotic phenotype. We therefore decided to primarily focus on the remaining components in this study. However, it is worth noting that in *C. elegans* a dynein light chain protein Dlc1 is involved in stabilizing the m6A writer Mett10 (Dorsett & Schedl, 2009). More work is needed to dissect the role of Dyn2 in the yeast MTC.

### Multifunctionality of the yeast MTC

Previous studies have pointed at m6A-dependent and m6A-independent roles played by Ime4 (Agarwala *et al*., 2012; Clancy *et al*., 2002). These conclusions were primarily guided by differences in phenotypes between *IME4* deletion and an *IME4* catalytic inactive mutant, both of which abolishing methylation but are associated with phenotypes of distinct severity. The nature of such m6A-independent roles had been so far unknown. Such a phenotype could be ‘MTC complex independent’, i.e. mediated via a completely new function of ime4, unrelated not only to its catalytic activity but also to its participation in the methylation-complex. Alternatively, such a phenotype could be MTC-dependent. The discovery of additional components of the yeast MTC allowed us to uncover that the substantially more severe phenotype of the *IME4* deletion mutant was shared also upon loss of other components of the yeast MTC, strongly suggestive of an MTC-dependent function. These results suggest that an intact MTC is critical for yeast meiosis, independent of its ability to methylate RNA. In this sense, these results are reminiscent of studies conducted on diverse proteins playing a role in modifying the ribosomal RNA (rRNA) or transfer RNA (tRNA), often revealing that catalytically inactive mutations display considerably less pronounced phenotypes than full deletions (Letoquart *et al*, 2014; Leulliot *et al*, 2008; Sharma *et al*, 2015; Sharma *et al*, 2013; White *et al*, 2008). In such cases, there was no moonlighting activity of the modifying enzyme reported, but instead the modifying enzyme played a scaffolding or chaperoning role, for which its catalytic activity was not essential.

m6A-independent functions have been reported for METTL3 and METTL14 of the mammalian MTC, however, to our knowledge no role for an inactive but intact MTC in mammals has been documented so far (Barbieri *et al*, 2017; Liu *et al*, 2021). In addition to the m6A dependent and independent roles of the yeast MTC in meiosis, Kar4 also has a separate function where it acts as a transcription factor in mating (Kurihara *et al*., 1996; Lahav *et al*., 2007). The functions in transcription and in the MTC of Kar4 are not linked since Kar4 does not stably associate with chromatin during meiosis.

At the time of preparation of this manuscript a related study showed that Ylg036w-Vir1 is important for m6A deposition and meiosis in yeast (Park *et al*, 2023a). Additionally, the same group showed in two different studies that Kar4 has an important role in controlling gene expression in meiosis, and elegantly identified separation of function mutants that affect either the mating or meiotic function of Kar4 (Park *et al*, 2023b; Park *et al*, 2023c). Both works support the findings described in our manuscript.

### Concluding remarks

In conclusion, we describe a new composition of the yeast MTC. We showed that yeast MTC components are considerably more conserved than previously described, and yet the yeast MTC is re-configured compared to mammalian and Drosophila MTCs. The yeast MTC exerts both m6A-dependent and m6A-independent functions that are both important for meiosis. Our findings expand the relevance of yeast as a model system for understanding the molecular mechanisms regulating m6A deposition, as well as other m6A-independent roles of MTCs.

## Methods

### Plasmids and yeast strains

For this study we used strains derived from the sporulation proficient SK1 background and experiments were carried out in diploid cells unless stated otherwise. Gene deletions and carboxy terminal tagging with mNeongreen, 3xV5, TEV Protein A, or AID were performed using one step gene replacement protocol described previously (Longtine *et al*, 1998; Tam & van Werven, 2020). Depletions using C-terminal auxin-inducible degron (AID) tag were performed as described by (Nishimura *et al*, 2009). A short version of the AID-tag, known as mini-AID, was used for depleting Slz1, Pab1 and Npl3 (Morawska & Ulrich, 2013). To induce depletion during meiosis we used copper inducible *Oryza sativa* TIR (*osTIR*) ubiquitin ligase under the control of the *CUP1* promoter (Varier *et al*., 2022). The strain genotypes are listed in Table S3, the plasmids used in Table S4, and the oligo sequences in Table S5.

### Growth conditions

Cells were grown in YPD (1.0% (w/v) yeast extract, 2.0% (w/v) peptone, 2.0% (w/v) glucose, and supplemented with uracil (2.5 mg/l) and adenine (1.25 mg/l)). To induce meiosis a standard protocol for sporulation was followed as described previously (Varier *et al*., 2022). In short, cells were grown until saturation for 24h in YPD, then grown in pre-sporulation medium BYTA (1.0% (w/v) yeast extract, 2.0% (w/v) bacto tryptone, 1.0% (w/v) potassium acetate, 50 mM potassium phthalate) for about 16 h, and shifted to sporulation medium (SPO) (0.3% (w/v) potassium acetate and 0.02% (w/v) raffinose)). All experiments were performed at 30°C in a shaker incubator at 300 rpm. For mating experiments, cells were grown to early log phase in YPD, when 1uM α-factor was added, and cells were incubated with the pheromone for 30 minutes before collection. To enable efficient depletion of mini-AID strains (Slz1, Vir1, Pab1 and Npl3), 1 mM of indole-3-acetic acid (IAA) and 50 µM of CuSO_4_ were added 2 hours after cells were shifted to SPO and samples collected at 4 hours in SPO. As a control, the same volume of dimethyl sulphoxide (DMSO) was added to the yeast culture.

### RNA extraction

RNA extraction was performed from yeast pellets using Acid Phenol:Chloroform pH 4.5 and Tris-EDTA-SDS (TES) buffer (0.01 M Tris-HCl pH 7.5, 0.01 M EDTA, 0.5% w/v SDS). Samples were further treated with rDNase (cat no 740.963, Macherey-Nagel) and column purified (cat no 740.948, Macherey-Nagel).

### RT-qPCR

For reverse transcription, ProtoScript II First Strand cDNA Synthesis Kit (New England BioLabs) was used and 500 ng of total RNA was provided as template in each reaction. qPCR reactions were prepared using Fast SYBR Green Master Mix (Thermo Fisher Scientific) and transcript levels were quantified from the cDNA on Quantstudio 7 Flex Real Time PCR instrument. Signals were normalised over *ACT1*.

### DAPI counting

Cells were collected from sporulation cultures, pelleted via centrifugation, and fixed in 80% (v/v) ethanol for a minimum of 2h at 4°C. The cells were resuspended in PBS with 1 µg/ml DAPI. The proportion of cells containing two or more nuclei were considered meiosis.

### ChIP and ChIP-seq

Chromatin immunoprecipitation (ChIP) was performed as described previously (Moretto *et al*, 2018). In short, cells were fixed in 1.0% v/v formaldehyde. Cells were lysed in FA lysis buffer (50 mM HEPES–KOH, pH 7.5, 150 mM NaCl, 1mM EDTA, 1% Triton X-100, 0.1% Na-deoxycholate, 0.1% SDS and protease cocktail inhibitor (complete mini EDTA-free, Roche)) and chromatin was sheared by sonication using a Bioruptor (Diagenode, 9 cycles of 30 s on/off). Extracts were incubated for 2 h at room temperature with anti-V5 agarose beads (Sigma), washed twice with FA lysis buffer, twice with wash buffer 1 (FA lysis buffer containing 0.5M NaCl), and twice with wash buffer 2 (10 mM Tris–HCl, pH 8.0, 0.25M LiCl, 1 mM EDTA, 0.5% NP-40, 0.5% Na-deoxycholate). Reverse cross-linking was done in 1% SDS-TE buffer + Ribonuclease A (10 ng/μl) (100 mM Tris pH 8.0, 10 mM EDTA, 1.0% v/v SDS) at 65°C overnight. Proteinase K treated samples were column purified.

qPCR was used for determining the association with AGA1 promoter, and HMR was used a background control. For ChIP-seq, libraries were prepared using the KAPA Hyperprep kit (Roche) according to the manufacturer’s protocol. Libraries were size-selected for DNA fragments between 150 nt and 700 nt using gel extraction. Purified libraries were further quantified and inspected on a Tapestation (Agilent Technologies) and sequenced on an Illumina HiSeq 2500 to an equivalent of 50 bases single-end reads, at a depth of approximately 16 million reads per library.

### ChIP-seq data analysis

For ChIP-seq data adapter trimming was performed with cutadapt (version 1.9.1) with parameters “--minimum-length=25 --quality-cutoff=20 -a AGATCGGAAGAGC,” while for Ndt80 ChIP-seq, the cutadapt parameters are “-a AGATCGGAAGAGC -- minimum-lengthC=C20.” BWA (version 0.5.9-r16) with default parameters was used to perform genome-wide mapping of the adapter-trimmed reads to the SK1 genome (Li *et al*, 2009; Martin, 2011). Duplicate marking was performed using the picard tool MarkDuplicates (version 2.1.1) (http://broadinstitute.github.io/picard). Further filtering was performed to exclude reads that were duplicates and ambiguously mapped.

The peak calling for ChIP-seq data was done by using “macs2 callpeak” in MACS2 with parameters “-g 12000000 -m 3 100 -B -q 0.05” (Liu, 2014). Based on the output of “macs2 callpeak,” the signal tracks were generated by the “macs2 bdgcmp” command with parameters “-m ppois” (Heinz *et al*, 2010; Liu, 2014; Quinlan & Hall, 2010).

### Western blotting and quantification

3.6 ODs of yeast cells were pelleted from cultures via centrifugation for protein expression analysis. Proteins were precipitated from whole cells via trichloroacetic acid (TCA). Cells were lysed with glass beads using a bead beater and lysis buffer (50 mM Tris, 1 mM EDTA, and 2.75 mM DTT). Proteins were denatured in 3X sample buffer (9% (w/v) SDS and 6% (v/v) β-mercaptoethanol) at 100°C and separated by SDS/PAGE (4-12% gels). A PVDF membrane was used for protein transfer. Blocking was performed using 5% (w/v) dry skimmed milk. Proteins were detected using an ECL detection kit or using infra-red fluorescent antibodies visualised with LiCor CLx. Samples were normalised using Hxk1 as a loading control. Images were taken using the Li-Cor (manufacturer) and analysed using the Image Studio Lite software.

For the chemical RNA interactome capture (RIC) experiments, 15 μg of total protein (input control) and 20% of eluates were resolved on 4-20% SDS polyacrylamide gels (SDS-PAGE) and transferred to nitrocellulose (NC) membranes (Cytiva life sciences, 10600003). Membranes were blocked in PBS-0.1% Tween-20 containing 5% skimmed milk, probed with designated antibodies and horseradish peroxidase (HRP)-coupled secondary antibodies, and developed with the Immobilon Western Chemiluminescent HRP Substrate (MerckMillipore, WBKLS0500). Blots were recorded with an Amersham A600 gel documentation system (Cytiva life sciences). The following antibodies were used: V5 tag monoclonal antibody (1:2,000; ThermoFisher, R96025), mouse anti-Pab1 (1:5000; Antibodies online, ABIN1580454), mouse anti-Act1 (1:2500; MP Biomedicals, 0869100), HRP-conjugated sheep anti-mouse IgG (1:5000; Cytiva, NA931V).

### Immunoprecipitation and mass spectrometry

Yeast cells containing the Mum2-TEV-ProA tagged protein and control cells were induced to undergo sporulation as previously described (Spedale *et al*, 2010). For each immunoprecipitation experiment we used 1250 OD units of yeast cells. Cells were resuspended in lysis Buffer E (10 mM HEPES-NaOH, pH 8.0, 150mM NaCl, 0.1% Tween-20, 10% glycerol, 1mM dithiothreitol (DTT)) supplemented with RNase inhibitors (HALT protease inhibitor), and snap frozen, and processed into a powder using a Spex SamplePrep Freezer/Mill Cryogenic Grinder. Cell powder was dissolved in 12 ml Lysis Buffer E supplemented with RNase inhibitors and rotated to mix for at least 30 minutes in 4°C. Large debris was removed by a short spin (3000g for 2 minutes, 4°C), and to clarify the lysate 45.000 rpm for 1 hour. All handling of the lysate was done at 4°C. For each sample, 250 μl Streptavidin M-280 Dynabeads (Thermo Fisher Scientific) coated with 30 μl Biotinylated Anti-Rabbit IgG (Vector Laboratories) was added. The yeast extracts were incubated with the beads for 2 hours at 4°C. For samples treated with RNase I, the protein concentration was determined with Bradford reagent (manufacturer) and 1 U RNase I per mg protein was added at the start of the incubation. Beads were collected using magnetic stands, washed once with lysis buffer E, and washed 3 times with Cleavage Buffer (10 mM Tris-HCl, pH 8.0, 150 mM NaCl, 0.5 mM EDTA, 0.1% Tween-20, 1 mM DTT). Finally, beads were resuspended in 49 μl cleavage buffer. Proteins were eluted by the addition of 1uL TEV Protease (New England Biolabs) and incubated for 2 hours at room temperature, gently shaking at 450 rpm.

Reduced and alkylated proteins were prepared by in-gel digesting overnight at 37°C using 100ng trypsin (Promega). The supernatant was dried by vacuum centrifugation and the samples were resuspended in 0.1% trifluoro acetic acid. 1-10 μl of acidified protein digest was loaded on an Ultimate 3000 nanoRSLC HPLC (Thermo Scientific) onto a 20mm x 75um Pepmap C18 trap column (Thermo Scientific) prior to elution via a 50cm x 75um EasySpray C18 column into a Lumos Tribrid Orbitrap mass spectrometer (Thermo Scientific). A 70-minute gradient of 6% - 40% B was used followed by washing and re-equilibration (A= 2%ACN, 0.1% formic acid; B= 80%ACN, 0.1% formic acid). The Orbitrap was operated in “Data Dependent Acquisition” mode followed by MS/MS in “TopS” mode using the vendor supplied “universal method” with default parameters. Raw files were processed using Maxquant (maxquant.org) and Perseus (maxquant.net/perseus) with a recent download of the UniProt *Saccharomyces cerevisiae* reference proteome database and a common contaminants database. A decoy database of reversed sequences was used to filter false positives, at a peptide false detection rate of 1%. T-tests were performed with a permutation-based FDR of 5% to address multiple hypothesis testing.

### Fluorescence microscopy and image quantification

Kar4-mNeongreen and Htb2-mCherry image acquisition was conducted using a Nikon Eclipse Ti inverted microscope. Exposure times were set as follows: 500ms GFP, and 50ms mCherry. An ORCA-FLASH 4.0 camera (Hamamatsu) and NIS-Elements AR software (Nikon) were used to collect images. Quantification of fluorescence signals was performed using ImageJ (Schindelin *et al*, 2012). ROIs were manually drawn around the periphery of each cell or around the nucleus. The mean intensity in each channel per cell was determined and used for the analysis. Values for nuclear protein localisation were derived via the division of nuclear / whole cell signal. For the analyses, 50 cells were quantified per sample.

### m6A-mRNA quantification by LC-MS/MS and m6A-ELISA

RNA was extracted and DNase treated as described above and polyA RNA isolated after purification twice with oligodT dynabeads (Ambion, 61005) according to manufacturer’s instructions. M6A LC-MS was described previously (Varier *et al*., 2022). In short, polyA selected RNA was digested with addition of reaction buffer (2.5mM ZnCl2, 25mM NaCl, 10mM NaAcetate), 2 units of Nuclease P1 (Sigma, N8630) and incubated for 4 h at 37 °C. Followed by 2 h with 100 mM ammonium bicarbonate, 1μl Alkaline Phosphatase (NEB, M0525). Formic Acid was added to the reaction mixture and filtered with a 1.5 ml microfuge tube. The sample was then injected (20 μl) and analysed by LC-MS/MS using a reverse phase liquid chromatography C18 column and a triple quadrupole mass analyser (Agilent 6470 or Thermo Scientific TSQ Quantiva) instrument in positive electrospray ionisation mode. Flow rate was at 0.2 ml/min and column temperature 25 °C with the following gradient: 2 min 98% eluent A (0.1% formic acid and 10 mM ammonium formate in water) and 2% eluent B (0.1% formic acid and 10 mM ammonium formate in MeOH), 75% A and 25% B up to 10 min, 20% A and 80% B up to 15 min, 98% A and 2% B up to 22.5 min. Nucleosides were quantified using a calibration curve of pure nucleosides standards.

For ELISA we either used the EpiQuik m6A RNA methylation quantification kit from EpiGentek (P-9005) according to manufacturer’s protocol or an m6A ELISA protocol previous described (Ensinck *et al*., 2023).

### Chemical crosslinking RNA-protein interactome capture

To determine Ime4 and Kar4 binding to mRNAs, we adapted an RNA-protein interactome protocol by using chemical crosslinking instead of UV crosslinking (Matia-Gonzalez *et al*, 2021; Na *et al*, 2021; Patton *et al*, 2020). In short, RNA-protein complexes were crosslinked chemically using 0.2% formaldehyde (ThermoFisher, 28908) for 10 min at 30 °C. Crosslinking was quenched by adding 2.5 M glycine to a final concentration 125 mM followed by incubation for 5 min in an orbital shaker at 30 °C. Cells were collected by centrifugation, washed twice with PBS, then flash-frozen in liquid nitrogen. Cells were ground in liquid nitrogen using pre-chilled mortar and pestle. Cell powder was resuspended in 3.5 ml of RIC lysis buffer (20 mM Tris-HCl pH 7.4, 600 mM LiCl, 0.5% LiDS, 2 mM EDTA, 5 mM DTT, 1% Triton X-100, 1.5× Halt™ Protease Inhibitor Cocktail (ThermoFisher, 78429), 1 mM AEBSF and 0.2 U SUPERase RNase inhibitor (Invitrogen, AM2696)). Lysates were clarified by centrifugation. Total protein quantification was estimated with Bradford assay.

RNA-protein complexes were isolated with oligo[dT)]_25_ magnetic beads (New England Biolabs, S1419S). Briefly, 350 µl of oligo[dT)]_25_-coupled beads were pre-equilibrated by washing them five times with 1 ml of RIC lysis buffer. 3.5 mg of total protein extracts from each culture were mixed with beads, vortexed briefly then incubated at room temperature for 15 min with gentle rotation in a rotator wheel. Beads were washed twice with 1 ml of wash buffer A (20 mM Tris-HCl pH 7.5, 600 mM LiCl, 0.2% LiDS, 0.5 mM EDTA, 0.1% Triton X100, 1× 1.5× Halt™ Protease Inhibitor Cocktail,1 mM AEBSF and 0.2 U SUPERase), and twice with 1 ml of wash buffer B (20 mM Tris-HCl pH 7.5, 600 mM LiCl, 0.5 mM EDTA, 1× 1.5× Halt™ Protease Inhibitor Cocktail,1 mM AEBSF and 0.2 U ml SUPERase) each for 15 seconds on ice. Beads were resuspended in 60 µl elution buffer (10 mM Tris-HCl pH 7.5, 10 U of RNase I, 4 µg RNase A) and incubated at 37 °C for 10 minutes. For mock elution control, beads were resuspended in elution buffer without RNases.

### Sequence Alignment

Whole proteome sequences were acquired from UniProt (UniProt, 2023). Global and local sequence alignment scores were determined using EMBOSS needle and matcher, respectively, with the emboss/6.6.0 package (Rice *et al*, 2000) (Madeira *et al*, 2019). Parameters employed for needle and water were EBLOSUM62 matrix with Gap_penalty: 10.0 and Extend_penalty: 0.5. Parameters for matcher were Gap_penalty: 14 and Extend_penalty: 4.

### Structural alignment

Structural predictions were downloaded from the AlphaFold database (Jumper *et al*, 2021; Varadi *et al*, 2022). Structural similarity was calculated using the TM-align algorithm (Zhang & Skolnick, 2005), using TM-align/20210224. The resultant TM-score was normalized by the length of the shared query protein (e.g. when comparing *YGL036W* to the whole Human genome, *YGL036W* length was used).

### m6A-seq2

m6A levels were determined by m6A-seq2, as previously described (Dierks *et al*, 2021; Schwartz *et al*., 2013). Briefly, RNA was poly-A selected twice using Dynabeads mRNA DIRECT kit, and fragmented to ∼150 bp fragments. 3’ RNA barcode adaptors were then ligated (see Table S6), and all samples were pooled for m6A-IP-based enrichment. 90% of the mRNA material was subjected to two rounds of immunoprecipitation using two distinct anti-m6A antibodies, as follows: The RNA was first incubated for four hours with protein G beads (invitrogen) coupled to a Synaptic Systems poly-clonal m6A antibody (cat. 7945). Next, it was incubated overnight with protein A beads (invitrogen) coupled to a Cell Signaling poly-clonal m6A antibody (cat. D9D9W). RNA from input and IP libraries were reverse transcribed and PCR amplified as previously described (Dierks *et al*., 2021). cDNA libraries were quantified using Qubit RNA HS kit (Life technologies) and Tapestation High Sensitivity D1000 ScreenTape (Agilent Technologies), pooled with a ratio of ⅓ Input to ⅔ IP samples, and sequenced.

### m6A-seq2 data analysis

Paired-end reads were first demultiplexed according to barcodes in Read2 position 4-10, using an in-house python script. Reads were then aligned using STAR/2.5.3a to the SK1 reference genome (Schwartz *et al*., 2013) with the following additional parameters (--bamRemoveDuplicatesType UniqueIdentical, --outSAMtype BAM SortedByCoordinate,--alignIntronMax 500, --alignEndsType Local). Read were allocated to genes using txtools version 0.0.7.4 according to a SK1 gene annotation table. m6A sample index was calculated as the ratio of reads coverage between IP and Input samples, in a 51 bases window around a set of 1308 previously well-defined m6A sites (Table S6). To ensure adequate quantification, only sites with an average of 5 reads per base in the input fraction were used.

### RNA-seq

RNA levels were quantified by sequencing 3’ RNA ends as previously described (Chapal *et al*, 2019). Briefly, RNA was reverse transcribed using oligo dT primers coupled with internal barcodes [tag-seq barcodes Supp table]. RNA-DNA hybrids were then pooled and tagmented using Tn5, adding adaptors to their 5’ end, followed by PCR amplification of the fragments, inserting Illumina barcodes and adaptors. A total of four biological replicates were sequenced for each mutant-timepoint combination, two of each were processed in each of two pools.

### RNA-seq data analysis

Paired-end reads were demultiplexed according to barcodes in Read2 position 1-6, using an in-house python script., Read1 only were then aligned using STAR/2.5.3a (Dobin *et al*, 2013), with the following additional parameters (--outSAMtype BAM Unsorted, --alignIntronMax 500,--alignEndsType Local, --outBAMsortingBinsN 300, --limitBAMsortRAM 1254135158) and then deduplicated using UMI-tools 1.0.1. Reads were allocated to a custom-built transcriptome annotation of 3’ UTRs +-50 bp, using txtools 0.0.7.4. Transcription end site (TES) coordinates for SK1 cells entering meiosis were previously described (Chia *et al*, 2017). For a subset of transcripts, the UTR length was used to determine the end of transcripts (Nagalakshmi *et al*, 2008). Transcripts 3’ ends were annotated as ending at the farthest TES from the coding sequence and starting either 600 bases upstream of the closest TES or at the 5’ UTR start. The TES coordinates used are listed in Table S7. Samples with more than two standard deviations below the mean mapped reads per sample, or with low gene coverage (below 5000 genes mapped) were not included in the analysis (this excluded 3 samples).

Counts were normalized using DESEQ2/1.34.0 (Love *et al*, 2014). PCA analysis was done using PCAtools with removeVar = 0.1 option. Differentially expressed genes were determined by comparing mutated strain samples with their WT counterparts at matching timepoints, with the thresholds of adjusted p-value of 0.01 and an absolute log2 (FoldChange) of 1.5. Meiosis gene clusters were taken from a previously described dataset (Chu *et al*., 1998). Table S2 contains the RNA-seq data normalized readcounts.

### Cloning, expression and purification of recombinant yeast MTC

Ime4, Mum2, Slz1, Kar4, and Vir1 genes were codon-optimized for *E. coli* expression, synthesized *de novo*, and cloned into pACEBac1 (Epoch Life Sciences). During gene synthesis, a StrepII tag (SII) and site for cleavage by 3C PreScission protease were inserted at the N-terminal domain of Ime4 (SII-3C-Ime4). pACEBac1 plasmids were amplified by PCR using the original biGBac primers and introduced into pBIG1a by Gibson assembly using a modified version of the biGBac system (Hill *et al*, 2019. The final construct carrying the five subunits was verified using full-plasmid sequencing. Bacmid DNA was isolated from E. coli DH10 EmBacY cells, as described {Bieniossek, 2008 #365; Weissmann *et al*, 2016). To make P1 virus, 6-well dishes were seeded with 1.0 × 10^6^ *Sf9* cells per well in 2.0 mL InsectExpress medium (Lonza). Cells were transfected with 20 mL of bacmid per well, using Cellfectin II reagent as described by the manufacturer (Gibco). The supernatant (P1 virus) was harvested 72 h post-transfection, adding 40 mL of newborn calf serum (Gibco) before storage at 4 °C. P2 (amplified P1) virus was prepared by infecting suspension cultures of *Sf9* cells at 5.0 x 10^5^/mL with 10% v/v P1 virus and incubating for 4-5 days (130 rpm, 27 °C). Cells were checked for fluorescence, harvested by centrifugation (1000 g, 5 min), and the supernatant collected and stored at 4 °C. For further viral amplification, P3 (amplified P2) was prepared infecting Sf9 cells at 5.0 x 10^5^ cells/mL with 10% v/v P2 virus and incubating for 48h before harvesting the virus. Large-scale expression cultures were then set up by infecting 2–6 L suspension cultures of *High5* insect cells at 5.0 × 10^5^/mL with 0.5% v/v P3 virus. Following incubation (130 rpm, 27 °C), cells were harvested 48 hours post-infection by centrifugation (1000 g, 20 min, 4 °C), and stored at −80 °C.

Cell pellets from 2 L *High5* cells were resuspended in 50 mM HEPES pH 7.9, 300 mM NaCl, 1 mM PMSF, 10% glycerol, 1 mM TCEP, supplemented with 5 U/mL benzonase and EDTA-free protease inhibitors, and lysed by homogenization. The lysate was cleared by centrifugation (12,000 rpm, 30 min, 4°C), 0.45 μm-filtered, and loaded into a 5 mL StrepTrapTM HP column (Cytiva) pre-equilibrated in 50 mM HEPES pH 7.9, 300 mM NaCl, 1 mM PMSF, 1 mM TCEP. Elution was performed using the pre-equilibration buffer supplemented with 50 mM biotin. The purified complex was used immediately for SDS-PAGE analysis, and the sample was stored at 4 °C.

### Statistical analyses

Details of statistical tests used, sample number, and number of independent experiments are included in the relevant figure legends.

## Data and software availability

The GEO accession numbers for the m6A-seq2, RNA-seq, and ChIP-seq data reported in this manuscript are: GSE193561 and GSE224836.

## Acknowledgements

We thank the members of the van Werven lab for critical reading of the manuscript. This research was funded in whole, or in part, by the Wellcome Trust (FC001203). For the purpose of Open Access, the author has applied a CC BY public copyright licence to any Author Accepted Manuscript version arising from this submission. This work was supported by the Francis Crick Institute (FC001203), which receives its core funding from Cancer Research UK (FC001203), the UK Medical Research Council (FC001203), and the Wellcome Trust (FC001203).

## Author contributions

I.E., A.M. S.S. and F.J.v.W. conceived the project. I.E., A.M., S.S. and F.J.v.W. designed the experiments. I.E. performed most wet lab experiments, and A.M. performed most computations analysis, m6A-seq2, and RNA-seq experiments. W.A. developed and performed the polyA PD experiments in Figure 6. M.L. and G.S. under supervision of A.C. performed the in vitro reconstitution experiment in Figure 4. T.S. and E.C. performed the m6A-LC-MS experiments in Figure 1 under supervision of M.R.. S.H. performed and analysed the IP-MS samples under the supervision of A.P.S. and M.S. H.P. analysed the ChIP-seq data in Figure 6. I.E., A.M. S.S. and F.J.v.W. wrote the manuscript with input from the other authors. F.J.v.W. and S.S. supervised the project and provided funding.

## Supplementary data

Figures S1-S7

**Table S1.** MS data

**Table S2.** RNA-seq data

**Table S3.** Genotypes of strains used

**Table S4.** Oligo nucleotide sequences used

**Table S5.** Plasmids used

**Table S6.** List of Annotated m6A sites used for m6Aseq analysis

**Table S7.** TES coordinates used for RNA-seq

## Figure Legends

**Figure S1. Identification of Mum2 interacting proteins** (**A**) Diploid cell harbouring Kar4 tagged with TEV-ProA (FW9308) or untagged control (FW1511) were induced to enter meiosis and sample was taken at 4 hours in SPO. Protein extracts were generated and incubated with ProA coated paramagnetic beads. Subsequently, TEV protease was used to elute Mum2 from the beads. Shown are volcano plots of Kar4-TEV-ProA compared to untagged control. Kar4 and Ygl036w, which were enriched in Mum2 IP-MS of Figure 2B, are labelled in blue. Subunits of the MIS complex are labelled in red. **(B)** Similar analysis as in A, except that during Mum2-TEV-ProA and untagged control IP extracts were either untreated or RNase treated. Shown are volcano plots of Mum2-TEV-ProA compared to untagged control were either untreated (left panel) or RNase treated (right panel). Kar4, Ygl036w, Dyn2 and Pab1, which were enriched in Mum2 IP-MS of Figure 2B, are labelled in blue. Subunits of the MIS complex are labelled in red.

**Figure S2. Functional characterization of Mum2 interactors** Depletion of the *NPL3-AID*, *PAB1-AID*, *SLZ1-AID*, alleles (FW10388, FW10389, FW10386). Cells were induced to enter meiosis. After 4 hours in SPO cells were treated with IAA and CuSO4 or mock treated. Samples were taken at the indicated time points. Western blot membranes were probed with anti-V5 antibodies and anti-Hxk1 antibodies as a loading control. A representative experiment is shown.

**Figure S3. Ylg036w-Vir1 and Slz1 have orthologues in humans** Overlay of Alpha-fold predicted structures of Ylg036c-Vir1 and human VIRMA. Human VIRMA is shown in blue and Yeast Ylg036c-Vir1 is shown in pink. Also indicated are the Vir motif and regions with strong sequence similarities. Same figure as 3C but enlarged.

**Figure S4. Topological and structural properties of the yeast MTC** (**A-E**). Ime4, Mum2, Vir1, Kar4, and Slz1 protein expression depends on the presence of MTC subunits. Same data as described in Figure 4A. Western blots were probed with anti-V5 antibodies and anti-Hxk1 antibodies as a loading control. Western blot signals were quantified and the signal for the control at 0 hours was set to 1. The mean signal for n=2 biological repeats is shown. (**F**) mRNA expression (normalized read counts) of *IME4*, *MUM2*, *VIR1*, *SLZ1* and *DYN2* mRNA in control, *ime4*Δ, *mum2*Δ, *slz1*Δ, *kar4*Δ, *vir1*Δ and *dyn2*Δ (FW1511, FW7030, FW6535, FW6504, FW8246, FW9307 and FW10442) as determined by RNA-seq (See figure 5 for details) at 0h, 2h and 4h in SPO.

**Figure S5. m6A dependent and independent functions of the MTC in meiosis A)** Cells induced to enter meiosis. Samples were taken at the indicated time points for DAPI staining. Cells were fixed, stained, and DAPI masses were counted for at least 100 cells per biological repeat. Cells with two or more DAPI masses were considered meiosis. Similar analysis as A, except that catalytic dead mutant of Ime4 was analysed (*IME4^CD^*). For the analysis we used an *ime4* deletion mutant strain that had either plasmid integrated harbouring *IME4^WT^* or *IME4^CD^* (FW1511, FW7030, FW8736, and FW8773). The mean and SEM of n=3 biological repeats are displayed. **(B)** m6A levels in WT, *ime4*Δ, *IME4^WT^* or *IME4^CD^* (FW1511, FW7030, FW8736, and FW8773). Samples were collected at 4 hours in SPO of m6A-MS analysis. The mean of n=3 biological replicates are shown. **(C)** Clustering analysis of the RNA-seq experiment described in Figure 5C. **(D)** Venn diagram showing the transcripts that significantly changed in the deletion mutants of the 2-hour time point. **(E)** Median gene expression fold change from timepoint 0 of a random group of mRNAs (n=471) during the time course across the different deletion mutants. For each transcript a Z-score was calculated with respected to the 0-hour timepoint. The mean of at least n=3 biological replicates shown, and error bars represent the standard error of the mean.

**Figure S6. Kar4 function during mating and meiosis different** (**A**) Kar4 localization in cells untreated or treated α-factor. Kar4 fused to mNeongreen (mNg) was used for the analysis. To determine nuclear Kar4-mNg signal, we used histone H2B fused to mCherry (H2B-mCh). For the analysis we used *MAT*a and *MAT*a/α cells (FW8615 and FW8646). Quantification of nuclear (left panel) and whole cell (right panel) mean signal for Kar4-mNg. At least n=50 cells were quantified for the analysis. (**B**) Similar analysis as A, except *MAT*a/α diploid cells entering meiosis were used for the analysis. Cells were induced to enter meiosis in SPO, and samples were taken at 0, 3, and 6 hours in SPO. At least n=50 cells were quantified for the analysis. (**C**) ChIP-seq of control, Kar4-V5, Ime4-V5, Mum2-V5, and Slz1-V5 *MAT*a/α diploid cells (FW1511, FW8216, FW6057, FW6500, and FW6502). ChIP-seq data for the *GIC2* and *FUS1* loci are shown. (**D**) Motif analysis of Kar4 binding sites identified in α factor treated Kar4-V5 ChIP-seq data. Motifs were identified using the Homer software package. **(E)** Onset of meiosis in WT and *ste12*Δ (FW1511, FW9017). Cells induced to enter meiosis. Samples were taken at the indicated time points for DAPI staining. Cells were fixed, stained, and DAPI masses were counted for at least 100 cells per biological repeat. Cells with two or more DAPI masses were considered meiosis. The mean and SEM of n=3 biological repeats are displayed. **(F)** Chemical RNA-protein interactome analysis for assessing Ime4 and Kar4 binding to RNA. For the analysis we used Ime4-V5 and Kar4-V5 cells (FW6057 and FW8633). In short, cells entering meiosis (4h SPO) were crosslinked with formaldehyde, polyA mRNAs were pulled down from protein extracts with oligo-dT coated magnetic beads, washed, samples were eluted with RNase, and assessed by western blotting. Membranes were probed for anti-V5 to detect Ime4. As a positive control, membranes were probed with anti-Pab1. As negative control sample extract treated with RNase was included. **(G)** Photo activatable crosslinking of Kar4. Control and Kar4 (FW1511 and FW8633) cell entering meiosis were incubated with 4thiouracil (4TU). Cells were UV-crosslinked, and Kar4 was immunoprecipitated from protein extracts, and western blot membranes for Kar4 with anti V5 antibodies. * is an unrelated sample.

## References

Agarwala SD, Blitzblau HG, Hochwagen A, Fink GR (2012) RNA methylation by the MIS complex regulates a cell fate decision in yeast. PLoS Genet 8: e1002732

Aymoz D, Sole C, Pierre JJ, Schmitt M, de Nadal E, Posas F, Pelet S (2018) Timing of gene expression in a cell-fate decision system. Mol Syst Biol 14: e8024

Balacco DL, Soller M (2019) The m(6)A Writer: Rise of a Machine for Growing Tasks. Biochemistry 58: 363–378

Barbieri I, Tzelepis K, Pandolfini L, Shi J, Millán-Zambrano G, Robson SC, Aspris D, Migliori V, Bannister AJ, Han N et al (2017) Promoter-bound METTL3 maintains myeloid leukaemia by m6A-dependent translation control. Nature 552: 126–131

Bawankar P, Lence T, Paolantoni C, Haussmann IU, Kazlauskiene M, Jacob D, Heidelberger JB, Richter FM, Nallasivan MP, Morin V et al (2021) Hakai is required for stabilization of core components of the m(6)A mRNA methylation machinery. Nat Commun 12: 3778

Bujnicki JM, Feder M, Radlinska M, Blumenthal RM (2002) Structure prediction and phylogenetic analysis of a functionally diverse family of proteins homologous to the MT-A70 subunit of the human mRNA:m(6)A methyltransferase. J Mol Evol 55: 431–444

Bushkin GG, Pincus D, Morgan JT, Richardson K, Lewis C, Chan SH, Bartel DP, Fink GR (2019) m(6)A modification of a 3’ UTR site reduces RME1 mRNA levels to promote meiosis. Nat Commun 10: 3414

Chapal M, Mintzer S, Brodsky S, Carmi M, Barkai N (2019) Resolving noise-control conflict by gene duplication. PLoS Biol 17: e3000289

Chia M, Li C, Marques S, Pelechano V, Luscombe NM, van Werven FJ (2021) High-resolution analysis of cell-state transitions in yeast suggests widespread transcriptional tuning by alternative starts. Genome Biol 22: 34

Chia M, Tresenrider A, Chen J, Spedale G, Jorgensen V, Unal E, van Werven FJ (2017) Transcription of a 5’ extended mRNA isoform directs dynamic chromatin changes and interference of a downstream promoter. Elife 6

Chu S, DeRisi J, Eisen M, Mulholland J, Botstein D, Brown PO, Herskowitz I (1998) The transcriptional program of sporulation in budding yeast. Science 282: 699–705

Clancy MJ, Shambaugh ME, Timpte CS, Bokar JA (2002) Induction of sporulation in Saccharomyces cerevisiae leads to the formation of N6-methyladenosine in mRNA: a potential mechanism for the activity of the IME4 gene. Nucleic Acids Res 30: 4509–4518

Dagan Y, Yesharim Y, Bonneau AR, Frankovits T, Schwartz S, Reddien PW, Wurtzel O (2022) m6A is required for resolving progenitor identity during planarian stem cell differentiation. EMBO J 41: e109895

Deutschbauer AM, Williams RM, Chu AM, Davis RW (2002) Parallel phenotypic analysis of sporulation and postgermination growth in Saccharomyces cerevisiae. Proc Natl Acad Sci U S A 99: 15530–15535

Dierks D, Garcia-Campos MA, Uzonyi A, Safra M, Edelheit S, Rossi A, Sideri T, Varier RA, Brandis A, Stelzer Y et al (2021) Multiplexed profiling facilitates robust m6A quantification at site, gene and sample resolution. Nat Methods 18: 1060–1067

Dobin A, Davis CA, Schlesinger F, Drenkow J, Zaleski C, Jha S, Batut P, Chaisson M, Gingeras TR (2013) STAR: ultrafast universal RNA-seq aligner. Bioinformatics 29: 15–21

Dominissini D, Moshitch-Moshkovitz S, Schwartz S, Salmon-Divon M, Ungar L, Osenberg S, Cesarkas K, Jacob-Hirsch J, Amariglio N, Kupiec M et al (2012) Topology of the human and mouse m6A RNA methylomes revealed by m6A-seq. Nature 485: 201–206

Dorsett M, Schedl T (2009) A role for dynein in the inhibition of germ cell proliferative fate. Mol Cell Biol 29: 6128–6139

Ensinck I, Sideri T, Modic M, Capitanchik C, Vivori C, Toolan-Kerr P, van Werven F (2023) m6A-ELISA, a simple method for quantifying N6-methyladenosine from mRNA populations. RNA

Enyenihi AH, Saunders WS (2003) Large-scale functional genomic analysis of sporulation and meiosis in Saccharomyces cerevisiae. Genetics 163: 47–54

Gammie AE, Stewart BG, Scott CF, Rose MD (1999) The two forms of karyogamy transcription factor Kar4p are regulated by differential initiation of transcription, translation, and protein turnover. Mol Cell Biol 19: 817–825

Haussmann IU, Bodi Z, Sanchez-Moran E, Mongan NP, Archer N, Fray RG, Soller M (2016) m(6)A potentiates Sxl alternative pre-mRNA splicing for robust Drosophila sex determination. Nature 540: 301–304

Heinz S, Benner C, Spann N, Bertolino E, Lin YC, Laslo P, Cheng JX, Murre C, Singh H, Glass CK (2010) Simple combinations of lineage-determining transcription factors prime cis-regulatory elements required for macrophage and B cell identities. Mol Cell 38: 576–589

Hill CH, Boreikaite V, Kumar A, Casanal A, Kubik P, Degliesposti G, Maslen S, Mariani A, von Loeffelholz O, Girbig M et al (2019) Activation of the Endonuclease that Defines mRNA 3’ Ends Requires Incorporation into an 8-Subunit Core Cleavage and Polyadenylation Factor Complex. Mol Cell 73: 1217–1231 e1211

Huang H, Weng H, Zhou K, Wu T, Zhao BS, Sun M, Jiang X, Wu X, Sun M, Guan J-l, et al (2019) Histone H3 trimethylation at lysine 36 guides m6A RNA modification co-transcriptionally. Nature 567: 414–419

Jiang X, Liu B, Nie Z, Duan L, Xiong Q, Jin Z, Yang C, Chen Y (2021) The role of m6A modification in the biological functions and diseases. Signal Transduct Target Ther 6: 74

Jumper J, Evans R, Pritzel A, Green T, Figurnov M, Ronneberger O, Tunyasuvunakool K, Bates R, Zidek A, Potapenko A et al (2021) Highly accurate protein structure prediction with AlphaFold. Nature 596: 583–589

Knuckles P, Lence T, Haussmann IU, Jacob D, Kreim N, Carl SH, Masiello I, Hares T, Villasenor R, Hess D et al (2018) Zc3h13/Flacc is required for adenosine methylation by bridging the mRNA-binding factor Rbm15/Spenito to the m(6)A machinery component Wtap/Fl(2)d. Genes Dev 32: 415–429

Kurihara LJ, Stewart BG, Gammie AE, Rose MD (1996) Kar4p, a karyogamy-specific component of the yeast pheromone response pathway. Mol Cell Biol 16: 3990-4002

Lahav R, Gammie A, Tavazoie S, Rose MD (2007) Role of transcription factor Kar4 in regulating downstream events in the Saccharomyces cerevisiae pheromone response pathway. Mol Cell Biol 27: 818–829

Lence T, Paolantoni C, Worpenberg L, Roignant JY (2019) Mechanistic insights into m(6)A RNA enzymes. Biochim Biophys Acta Gene Regul Mech 1862: 222–229

Letoquart J, Huvelle E, Wacheul L, Bourgeois G, Zorbas C, Graille M, Heurgue-Hamard V, Lafontaine DL (2014) Structural and functional studies of Bud23-Trm112 reveal 18S rRNA N7-G1575 methylation occurs on late 40S precursor ribosomes. Proc Natl Acad Sci U S A 111: E5518–5526

Leulliot N, Bohnsack MT, Graille M, Tollervey D, Van Tilbeurgh H (2008) The yeast ribosome synthesis factor Emg1 is a novel member of the superfamily of alpha/beta knot fold methyltransferases. Nucleic Acids Res 36: 629–639

Li H, Handsaker B, Wysoker A, Fennell T, Ruan J, Homer N, Marth G, Abecasis G, Durbin R, Genome Project Data Processing S (2009) The Sequence Alignment/Map format and SAMtools. Bioinformatics 25: 2078–2079

Liu J, Yue Y, Han D, Wang X, Fu Y, Zhang L, Jia G, Yu M, Lu Z, Deng X et al (2014) A METTL3-METTL14 complex mediates mammalian nuclear RNA N6-adenosine methylation. Nat Chem Biol 10: 93–95

Liu P, Li F, Lin J, Fukumoto T, Nacarelli T, Hao X, Kossenkov AV, Simon MC, Zhang R (2021) m(6)A-independent genome-wide METTL3 and METTL14 redistribution drives the senescence-associated secretory phenotype. Nat Cell Biol 23: 355-365

Liu T (2014) Use model-based Analysis of ChIP-Seq (MACS) to analyze short reads generated by sequencing protein-DNA interactions in embryonic stem cells. Methods Mol Biol 1150: 81–95

Longtine MS, McKenzie A, 3rd, Demarini DJ, Shah NG, Wach A, Brachat A, Philippsen P, Pringle JR (1998) Additional modules for versatile and economical PCR-based gene deletion and modification in Saccharomyces cerevisiae. Yeast 14: 953-961

Love MI, Huber W, Anders S (2014) Moderated estimation of fold change and dispersion for RNA-seq data with DESeq2. Genome Biol 15: 550

Madeira F, Park YM, Lee J, Buso N, Gur T, Madhusoodanan N, Basutkar P, Tivey ARN, Potter SC, Finn RD et al (2019) The EMBL-EBI search and sequence analysis tools APIs in 2019. Nucleic Acids Res 47: W636–W641

Martin M (2011) Cutadapt removes adapter sequences from high-throughput sequencing reads. EMBnetjournal 17: 10

Matia-Gonzalez AM, Jabre I, Gerber AP (2021) Biochemical approach for isolation of polyadenylated RNAs with bound proteins from yeast. STAR Protoc 2: 100929

Meyer KD, Saletore Y, Zumbo P, Elemento O, Mason CE, Jaffrey SR (2012) Comprehensive analysis of mRNA methylation reveals enrichment in 31 UTRs and near stop codons. Cell 149: 1635–1646

Morawska M, Ulrich HD (2013) An expanded tool kit for the auxin-inducible degron system in budding yeast. Yeast 30: 341-351

Moretto F, Wood NE, Kelly G, Doncic A, van Werven FJ (2018) A regulatory circuit of two lncRNAs and a master regulator directs cell fate in yeast. Nat Commun 9: 780

Murakami S, Jaffrey SR (2022) Hidden codes in mRNA: Control of gene expression by m(6)A. Mol Cell 82: 2236–2251

Na Y, Kim H, Choi Y, Shin S, Jung JH, Kwon SC, Kim VN, Kim JS (2021) FAX-RIC enables robust profiling of dynamic RNP complex formation in multicellular organisms in vivo. Nucleic Acids Res 49: e28

Nagalakshmi U, Wang Z, Waern K, Shou C, Raha D, Gerstein M, Snyder M (2008) The transcriptional landscape of the yeast genome defined by RNA sequencing. Science 320: 1344-1349

Nishimura K, Fukagawa T, Takisawa H, Kakimoto T, Kanemaki M (2009) An auxin-based degron system for the rapid depletion of proteins in nonplant cells. Nat Methods 6: 917–922

Park ZM, Belnap E, Remillard M, Rose MD (2023a) Vir1p, the yeast homolog of virilizer, is required for mRNA m6A methylation and meiosis. Genetics 224

Park ZM, Remillard M, Rose MD (2023b) Kar4 is Required for the Normal Pattern of Meiotic Gene Expression. bioRxiv

Park ZM, Sporer A, Kraft K, Lum K, Blackman E, Belnap E, Yellman C, Rose MD (2023c) Kar4, the Yeast Homolog of METTL14, is Required for mRNA m (6) A Methylation and Meiosis. bioRxiv

Patil DP, Chen CK, Pickering BF, Chow A, Jackson C, Guttman M, Jaffrey SR (2016) m(6)A RNA methylation promotes XIST-mediated transcriptional repression. Nature 537: 369–373

Patton RD, Sanjeev M, Woodward LA, Mabin JW, Bundschuh R, Singh G (2020) Chemical crosslinking enhances RNA immunoprecipitation for efficient identification of binding sites of proteins that photo-crosslink poorly with RNA. RNA 26: 1216–1233

Ping X-l, Sun B-f, Wang L, Xiao W, Yang X, Wang W-j, Adhikari S, Shi Y, Lv Y, Chen Y-s, et al (2014) Mammalian WTAP is a regulatory subunit of the RNA N6-methyladenosine methyltransferase. Cell Research 24: 177–189

Quinlan AR, Hall IM (2010) BEDTools: a flexible suite of utilities for comparing genomic features. Bioinformatics 26: 841–842

Rice P, Longden I, Bleasby A (2000) EMBOSS: the European Molecular Biology Open Software Suite. Trends Genet 16: 276–277

Ruzicka K, Zhang M, Campilho A, Bodi Z, Kashif M, Saleh M, Eeckhout D, El-Showk S, Li H, Zhong S et al (2017) Identification of factors required for m(6) A mRNA methylation in Arabidopsis reveals a role for the conserved E3 ubiquitin ligase HAKAI. New Phytol 215: 157–172

Schindelin J, Arganda-Carreras I, Frise E, Kaynig V, Longair M, Pietzsch T, Preibisch S, Rueden C, Saalfeld S, Schmid B et al (2012) Fiji: an open-source platform for biological-image analysis. Nat Methods 9: 676-682

Schwartz S, Agarwala SD, Mumbach MR, Jovanovic M, Mertins P, Shishkin A, Tabach Y, Mikkelsen TS, Satija R, Ruvkun G et al (2013) High-resolution mapping reveals a conserved, widespread, dynamic mRNA methylation program in yeast meiosis. Cell 155: 1409–1421

Schwartz S, Mumbach MR, Jovanovic M, Wang T, Maciag K, Bushkin GG, Mertins P, Ter-Ovanesyan D, Habib N, Cacchiarelli D et al (2014) Perturbation of m6A writers reveals two distinct classes of mRNA methylation at internal and 5’ sites. Cell Rep 8: 284–296

Scutenaire J, Plassard D, Matelot M, Villa T, Zumsteg J, Libri D, Seraphin B (2022) The S. cerevisiae m6A-reader Pho92 promotes timely meiotic recombination by controlling key methylated transcripts. Nucleic Acids Res

Shah JC, Clancy MJ (1992) IME4, a gene that mediates MAT and nutritional control of meiosis in Saccharomyces cerevisiae. Mol Cell Biol 12: 1078–1086

Sharma S, Langhendries JL, Watzinger P, Kotter P, Entian KD, Lafontaine DL (2015) Yeast Kre33 and human NAT10 are conserved 18S rRNA cytosine acetyltransferases that modify tRNAs assisted by the adaptor Tan1/THUMPD1. Nucleic Acids Res 43: 2242–2258

Sharma S, Yang J, Watzinger P, Kotter P, Entian KD (2013) Yeast Nop2 and Rcm1 methylate C2870 and C2278 of the 25S rRNA, respectively. Nucleic Acids Res 41: 9062–9076

Sledz P, Jinek M (2016) Structural insights into the molecular mechanism of the m(6)A writer complex. Elife 5

Spedale G, Mischerikow N, Heck AJ, Timmers HT, Pijnappel WW (2010) Identification of Pep4p as the protease responsible for formation of the SAGA-related SLIK protein complex. J Biol Chem 285: 22793–22799

Su S, Li S, Deng T, Gao M, Yin Y, Wu B, Peng C, Liu J, Ma J, Zhang K (2022) Cryo-EM structures of human m(6)A writer complexes. Cell Res 32: 982–994

Tam J, van Werven FJ (2020) Regulated repression governs the cell fate promoter controlling yeast meiosis. Nat Commun 11: 2271

UniProt C (2023) UniProt: the Universal Protein Knowledgebase in 2023. Nucleic Acids Res 51: D523–D531

Uzonyi A, Dierks D, Nir R, Kwon OS, Toth U, Barbosa I, Burel C, Brandis A, Rossmanith W, Le Hir H et al (2023) Exclusion of m6A from splice-site proximal regions by the exon junction complex dictates m6A topologies and mRNA stability. Mol Cell 83: 237–251 e237

Varadi M, Anyango S, Deshpande M, Nair S, Natassia C, Yordanova G, Yuan D, Stroe O, Wood G, Laydon A et al (2022) AlphaFold Protein Structure Database: massively expanding the structural coverage of protein-sequence space with high-accuracy models. Nucleic Acids Res 50: D439–D444

Varier RA, Sideri T, Capitanchik C, Manova Z, Calvani E, Rossi A, Edupuganti RR, Ensinck I, Chan VWC, Patel H et al (2022) N6-methyladenosine (m6A) reader Pho92 is recruited co-transcriptionally and couples translation to mRNA decay to promote meiotic fitness in yeast. Elife 11

Wang P, Doxtader KA, Nam Y (2016) Structural Basis for Cooperative Function of Mettl3 and Mettl14 Methyltransferases. Mol Cell 63: 306–317

Wang Y, Li Y, Toth JI, Petroski MD, Zhang Z, Zhao JC (2014) N6-methyladenosine modification destabilizes developmental regulators in embryonic stem cells. Nat Cell Biol 16: 191–198

Wang Y, Zhang L, Ren H, Ma L, Guo J, Mao D, Lu Z, Lu L, Yan D (2021) Role of Hakai in m(6)A modification pathway in Drosophila. Nat Commun 12: 2159

Weissmann F, Petzold G, VanderLinden R, Huis In ’t Veld PJ, Brown NG, Lampert F, Westermann S, Stark H, Schulman BA, Peters JM (2016) biGBac enables rapid gene assembly for the expression of large multisubunit protein complexes. Proc Natl Acad Sci U S A 113: E2564-2569

Wen J, Lv R, Ma H, Shen H, He C, Wang J, Jiao F, Liu H, Yang P, Tan L et al (2018) Zc3h13 Regulates Nuclear RNA m(6)A Methylation and Mouse Embryonic Stem Cell Self-Renewal. Mol Cell 69: 1028–1038 e1026

White J, Li Z, Sardana R, Bujnicki JM, Marcotte EM, Johnson AW (2008) Bud23 methylates G1575 of 18S rRNA and is required for efficient nuclear export of pre-40S subunits. Mol Cell Biol 28: 3151–3161

Yue Y, Liu J, Cui X, Cao J, Luo G, Zhang Z, Cheng T, Gao M, Shu X, Ma H et al (2018a) VIRMA mediates preferential m6A mRNA methylation in 31 UTR and near stop codon and associates with alternative polyadenylation. Cell Discovery 4

Yue Y, Liu J, Cui X, Cao J, Luo G, Zhang Z, Cheng T, Gao M, Shu X, Ma H et al (2018b) VIRMA mediates preferential m(6)A mRNA methylation in 3’UTR and near stop codon and associates with alternative polyadenylation. Cell Discov 4: 10

Zaccara S, Ries RJ, Jaffrey SR (2019) Reading, writing and erasing mRNA methylation. Nat Rev Mol Cell Biol 20: 608–624

Zhang M, Bodi Z, Mackinnon K, Zhong S, Archer N, Mongan NP, Simpson GG, Fray RG (2022) Two zinc finger proteins with functions in m(6)A writing interact with HAKAI. Nat Commun 13: 1127

Zhang Y, Skolnick J (2005) TM-align: a protein structure alignment algorithm based on the TM-score. Nucleic Acids Res 33: 2302–2309

