## Supplementary material for "The yeast RNA methylation complex consists of conserved yet reconfigured components with m6A-dependent and independent roles": Table S4

**Table S4. Oligo nucleotide sequences used.**

**Name	Sequence	reference**

IE34 pAGA1 ChIP qPCR fw	AGGGTACCTGTCACATATATTCTCA

IE35 pAGA1 ChIP qPCR rv	ATTATGTTACAGCCGCGTTTTG

HMR1F_RT	acgatccccgtccaagttatg

HMR1R_RT	cttcaaaggagtcttaatttccctg
