## Supplementary material for "The yeast RNA methylation complex consists of conserved yet reconfigured components with m6A-dependent and independent roles": Table S5

**Table S5. Plasmids used.**

**nr	name	reference**

p227	pWG444 NAT

p17	pFA6a-kanMX6

p147	pFA6a.URA3Mx6

p78	pFA6a-TEV-ProA-KanMX6

p255	pL264 KAN -3v5

p386	pCA13-mNeongreen(Yeast Optimized)-NAT

p491	3Pk-miniAID-kanMX

p782	Ime4-v5 TRP wt integration plasmid

p783	Ime4-v5 TRP catalytic dead integration plasmid
