## Supplementary figures and images for "The yeast RNA methylation complex consists of conserved yet reconfigured components with m6A-dependent and independent roles"

### Figure S1-S6

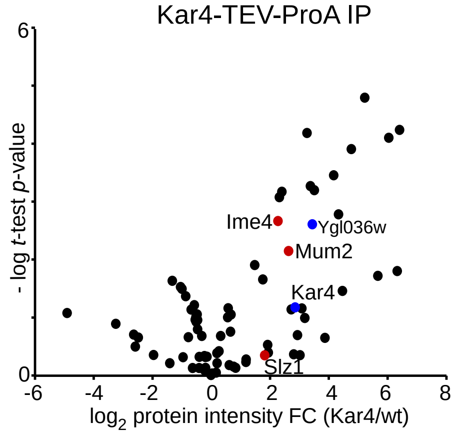

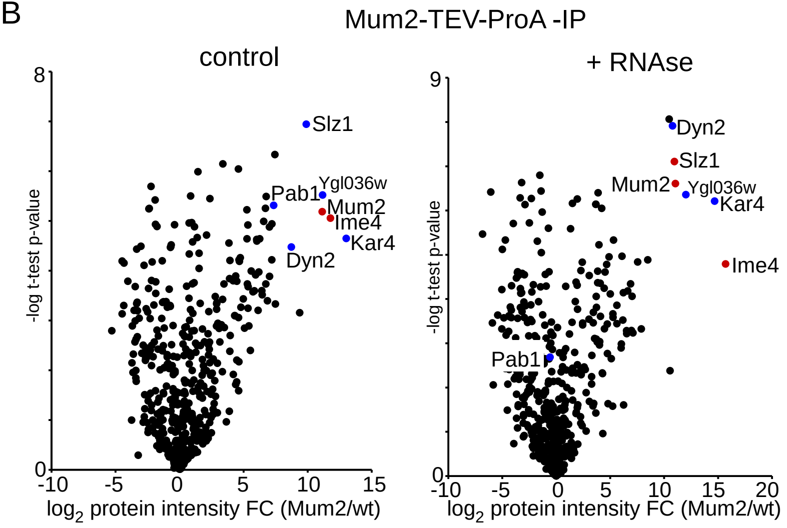

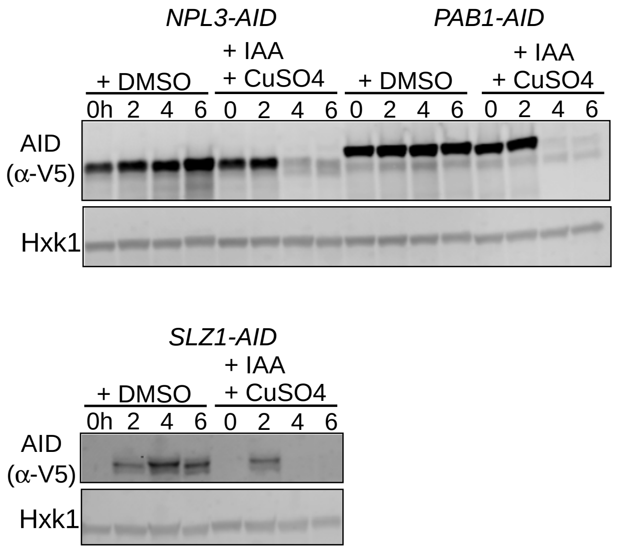

Figure S3

#### Ygl036w-Vir1 vs VIRMA

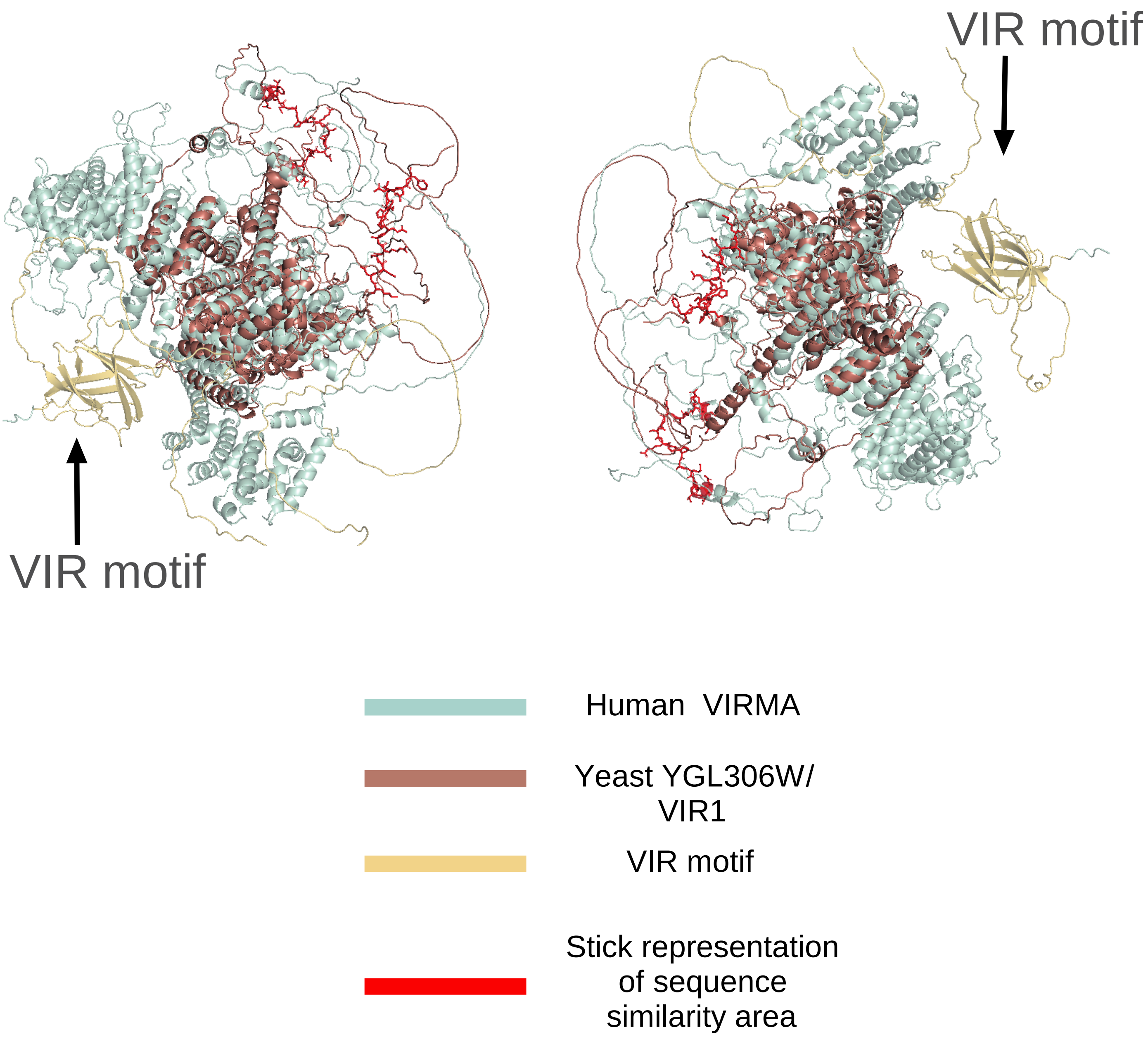

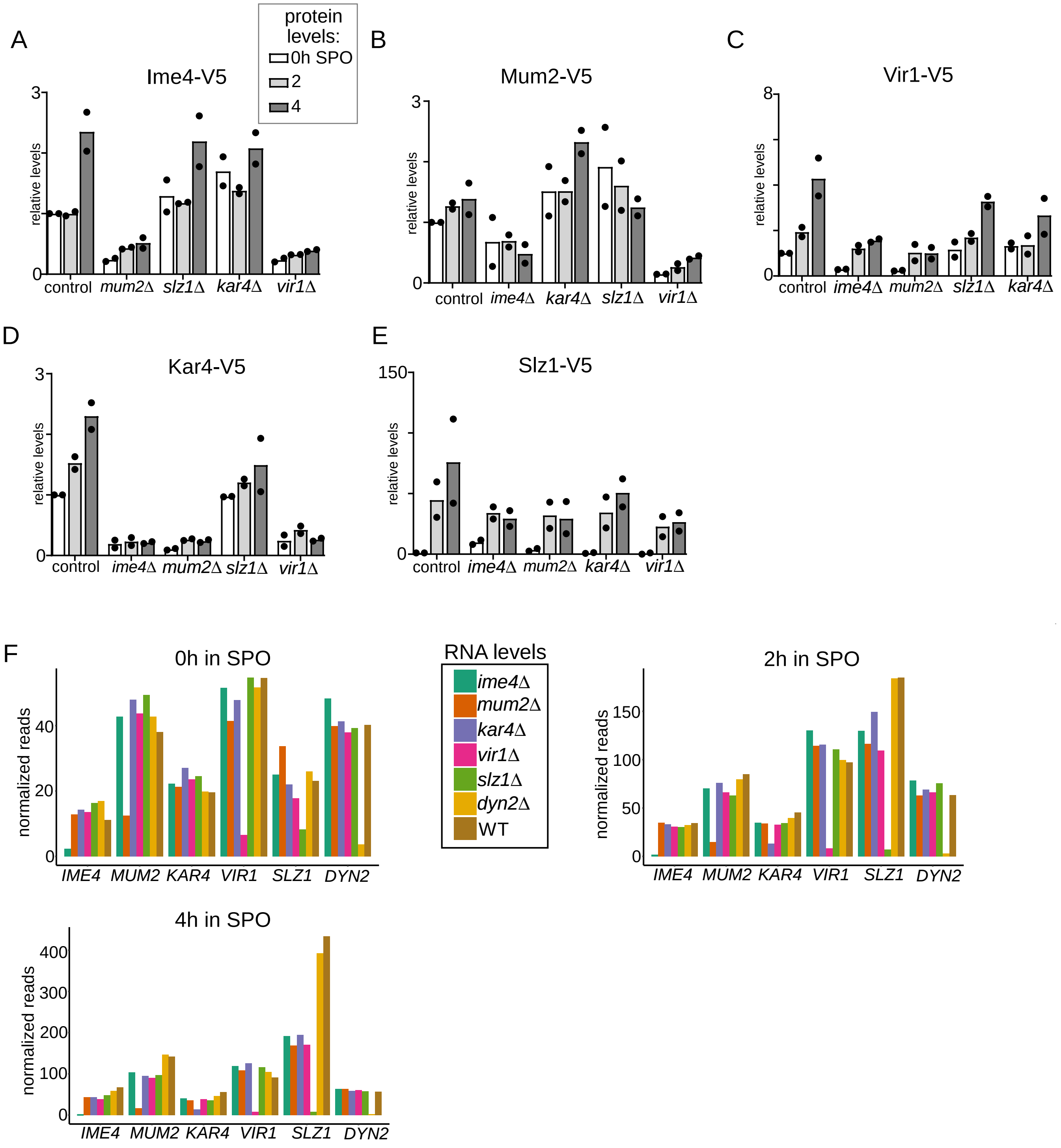

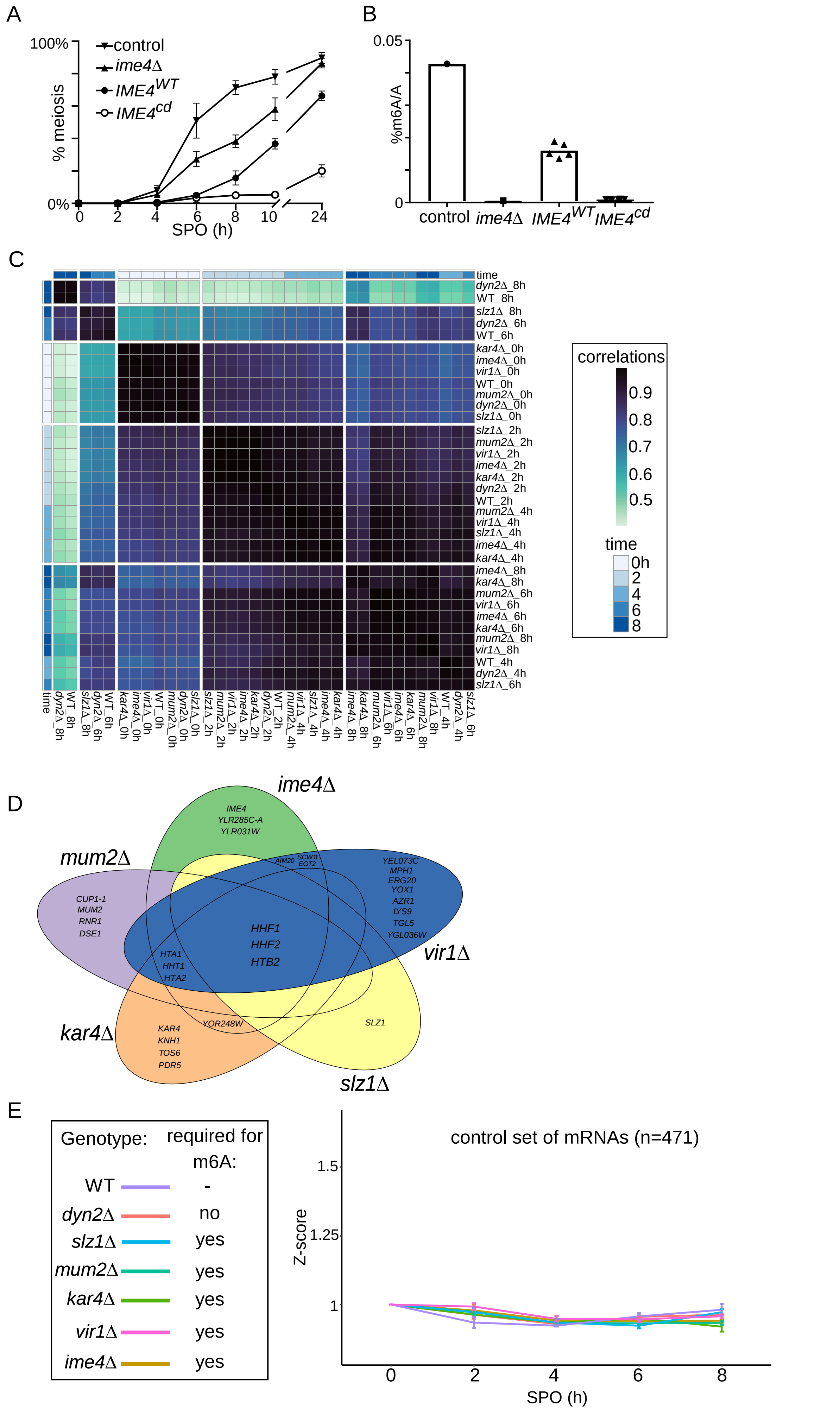

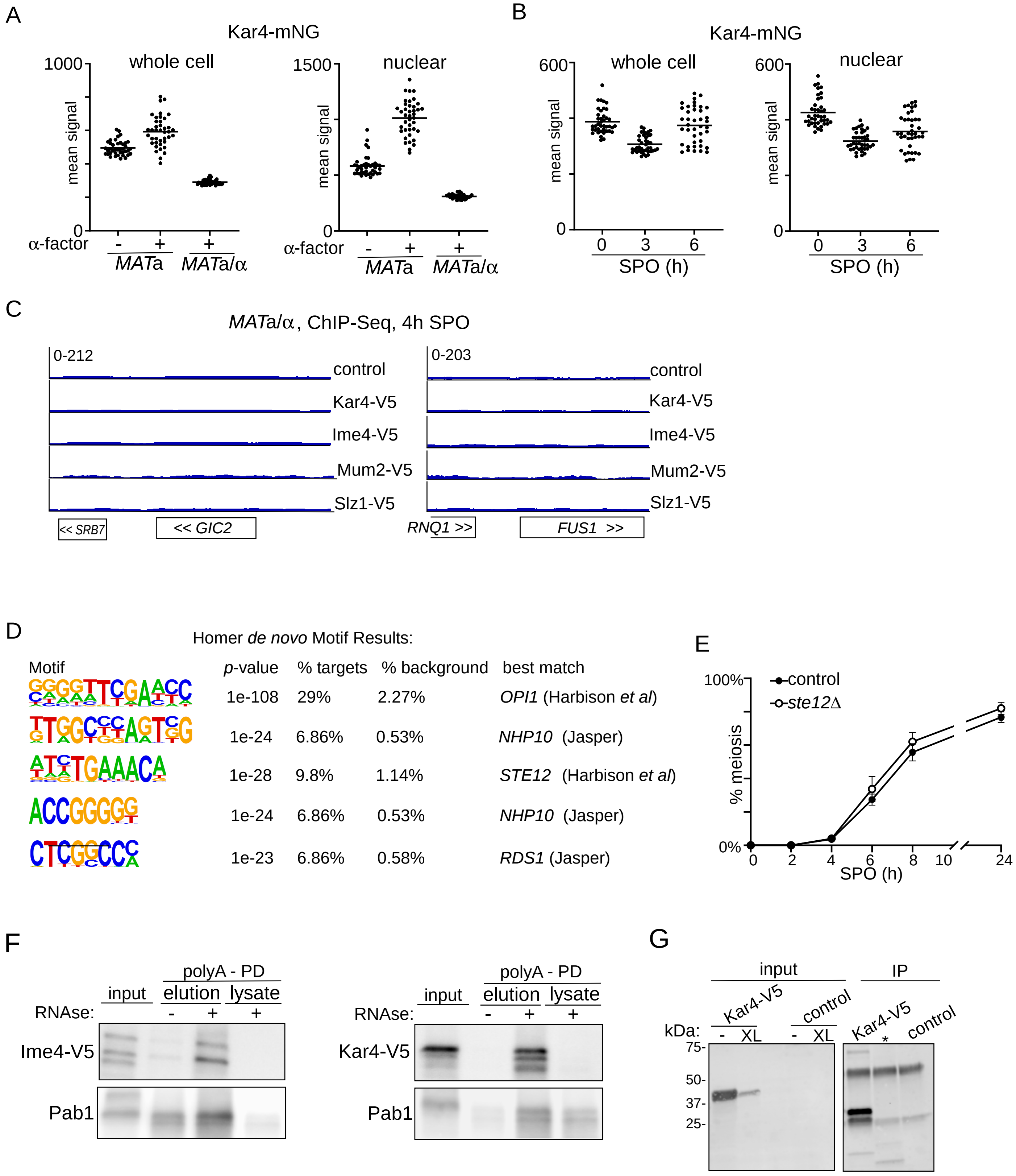
